# Inflammatory proteostasis reprogramming creates a lysosomal bottleneck for microglial aggregate clearance

**DOI:** 10.64898/2026.11.02.763615

**Authors:** Frederik Eikens, Lena Wischhof, Anna Henrich, Lena Erlebach, Annett Halle, Hannah Scheiblich

## Abstract

Efficient clearance of extracellular protein aggregates by microglia is essential for maintaining brain homeostasis and limiting the accumulation of pathological proteins during neurodegeneration. However, how inflammatory activation affects the ability of microglia to internalize and degrade these aggregates remains incompletely understood. Here, we show that lipopolysaccharide (LPS)-induced inflammatory activation impairs the degradation of extracellularly-acquired α-synuclein (α-Syn) and Tau aggregates while largely preserving their uptake. Mechanistically, Toll-like receptor 4 (TLR4) activation induces translational reprogramming and increases intracellular proteostatic demand, while inhibition of protein synthesis restores α-Syn degradation without preventing inflammatory Nuclear factor kappa-light-chain-enhancer of activated B cells (NF-κB) activation. Despite largely preserved canonical autophagic flux, inflammatory stimulation is associated with reduced expression of lysosomal proteins, including cathepsin D, suggesting insufficient lysosomal adaptation to increased intracellular demand. Consistent with this model, pharmacological activation of transcription factor EB (TFEB) restores aggregate degradation, indicating that the impairment can be overcome by enhancing lysosomal function. Supporting its potential relevance to human disease, microglial α-Syn deposits in brain tissue from patients with dementia with Lewy bodies show limited spatial association with cathepsin D-positive structures. Together, our findings identify a demand-driven lysosomal bottleneck through which inflammatory activation compromises aggregate degradation without substantially affecting uptake, highlighting lysosomal adaptation as a critical determinant of microglial aggregate clearance during neuroinflammation.

## Introduction

The accumulation and intercellular propagation of misfolded α-synuclein (α-Syn) are central pathological features of Parkinson’s disease (PD) and related synucleinopathies^1^. Under physiological conditions, α-Syn is an intrinsically disordered neuronal protein that regulates synaptic vesicle dynamics and neurotransmission^2^. During disease progression, however, α-Syn undergoes conformational transitions that promote its assembly into β-sheet-rich fibrils^3^, which template the recruitment of soluble monomers and ultimately form Lewy bodies, the pathological hallmark of PD. Importantly, α-Syn pathology is not restricted to the intracellular space. Misfolded α-Syn is continuously released from neurons and taken up by neighbouring cells, where it seeds further aggregation and contributes to the stereotypical spread of pathology across interconnected brain regions^1,4^. Efficient clearance of extracellular α-Syn is therefore likely to be an important determinant of α-Syn aggregate persistence and propagation in disease.

Microglia, the resident immune cells of the central nervous system, are the key regulators of extracellular proteostasis. In addition to orchestrating innate immune responses, they continuously survey the brain parenchyma and internalize extracellular material, including protein aggregates, cellular debris, and dying cells to maintain tissue homeostasis^5^. Microglia are particularly efficient at internalizing and degrading extracellular α-Syn aggregates, both *in vitro* and *in vivo*^6–8^, and this clearance capacity declines with age^7^. Importantly, microglial α-Syn clearance is sensitive to the inflammatory state of the cell. Lipopolysaccharide (LPS)-driven Toll-like receptor 4 (TLR4) activation has previously been shown to slow degradation of already internalized α-Syn aggregates^6^ and is increasingly discussed as a disease-relevant inflammatory stimulus in PD, as gut dysbiosis and increased intestinal permeability may enhance systemic endotoxin exposure and promote peripheral and central inflammation^9^. Other inflammatory paradigms, such as Interferon-γ (IFN-γ) stimulation of human induced pluripotent stem cells (iPSC)-derived microglia, have been shown to suppress α-Syn fibril uptake and disrupt subsequent intracellular processing^10^. Together, these observations suggest that inflammatory activation can affect distinct stages of microglial aggregate clearance, from initial internalization to subsequent degradation. However, changes in phagocytic uptake do not necessarily reflect the capacity of microglia to efficiently eliminate internalized material. During neurodegeneration, microglia face competing functional demands. They are required to rapidly mount inflammatory effector programs while simultaneously preserving their homeostatic capacity to eliminate extracellular pathogenic proteins. Although chronic neuroinflammation is increasingly recognized as a major contributor to PD progression^11^, how defined inflammatory programs such as TLR4 activation reconfigure the intracellular proteostatic machinery that supports extracellular aggregate degradation remains poorly understood.

Following internalization, extracellular cargo is generally routed through endolysosomal compartments toward degradation^12^. α-Syn pre-formed fibrils (PFFs) can undergo rapid macropinocytic uptake and lysosomal targeting^13^, although the intracellular route followed by α-Syn appears to depend on its form and mode of uptake. In particular, neuron-released α-Syn has been reported to engage TLR4-NF-κB-p62-dependent selective autophagy in microglia^14^. In parallel, damaged proteins, protein aggregates, and other intracellular components can be delivered to lysosomes through autophagy^15,16^. Although these pathways originate from distinct cellular compartments and trafficking routes, both ultimately converge at the lysosome^15,17^, making lysosomal function the common endpoint of intracellular protein quality control and extracellular aggregate degradation. Importantly, inflammatory activation profoundly increases the intracellular biosynthetic and proteostatic demand placed on immune cells by inducing rapid synthesis of cytokines, chemokines, antimicrobial proteins, and other inflammatory mediators. Whether these intracellular demands influence the ability of microglia to simultaneously clear extracellular protein aggregates remains unknown. More specifically, it remains unclear how the lysosomal system accommodates increased intracellular proteostatic demand while continuing to support efficient endolysosomal degradation of extracellular cargo. This raises the possibility that inflammation may compromise extracellular aggregate clearance even when the initial uptake of pathological proteins remains intact. We therefore reasoned that inflammatory activation might not simply impair lysosomal degradation *per se*, but instead disrupt the balance between intracellular proteostatic demand and the adaptive capacity of the lysosomal system.

To address this question, we systematically investigated how inflammatory activation affects the uptake and degradation of extracellular α-Syn in primary microglia. We demonstrate that inflammatory activation impairs degradation of extracellular α-Syn without substantially affecting its uptake. This reveals a functional uncoupling between aggregate internalization and degradation under inflammatory conditions. Mechanistically, inflammatory activation increases intracellular proteostatic demand without a corresponding increase in lysosomal capacity, creating a demand-driven lysosomal bottleneck that selectively limits extracellular aggregate degradation despite largely preserved canonical autophagic flux. This impairment extends to Tau aggregates, indicating that inflammation-induced impairment of extracellular aggregate degradation is not restricted to α-Syn. Restoration of lysosomal biogenesis is sufficient to overcome this degradative defect, identifying lysosomal adaptation as a critical determinant of microglial proteostasis during inflammation. Together, these findings highlight the importance of degradative capacity, rather than phagocytic uptake alone, in determining the efficiency of microglial aggregate clearance during neuroinflammation.

## Results

### Inflammatory activation induces broad proteostasis reprogramming in microglia

To assess how inflammatory activation changes the microglial proteome, we performed proteomics in primary mouse microglia after 24 h LPS stimulation (10 ng/ml) (Figure 1A). Principal component analysis revealed a clear separation between control and LPS-treated samples (Figure 1B). Overall, 3478 (1903 up, 1575 down) of 6742 detected proteins were differentially abundant, corresponding to just over half of the detected proteome (Figures 1C-D). KEGG pathway enrichment analysis revealed that downregulated proteins were predominantly associated with the tricarboxylic acid (TCA) cycle and oxidative phosphorylation, whereas upregulated proteins were mainly enriched in pathways related to inflammation, inflammatory signaling, and glycolysis (Figure 1E). Considering that LPS stimulation altered the abundance of nearly half of the detected proteome, we next asked how these broad changes affected the cellular proteostasis network. Module score analysis showed increased abundance of proteins associated with inflammatory signaling (Figure 1F), translation (Figure 1G), and proteasomal degradation (Figures 1H). In contrast, lysosome-associated proteins were strongly reduced (Figure 1I). Together, these data show that LPS does not affect all proteostatic pathways in the same way. While inflammatory signaling, protein synthesis, and proteasome-associated pathways increase, the lysosomal program is reduced (Figure 1J). This raised the question of whether inflammatory activation affects the ability of microglia to process extracellular cargo.

**Figure 1.**
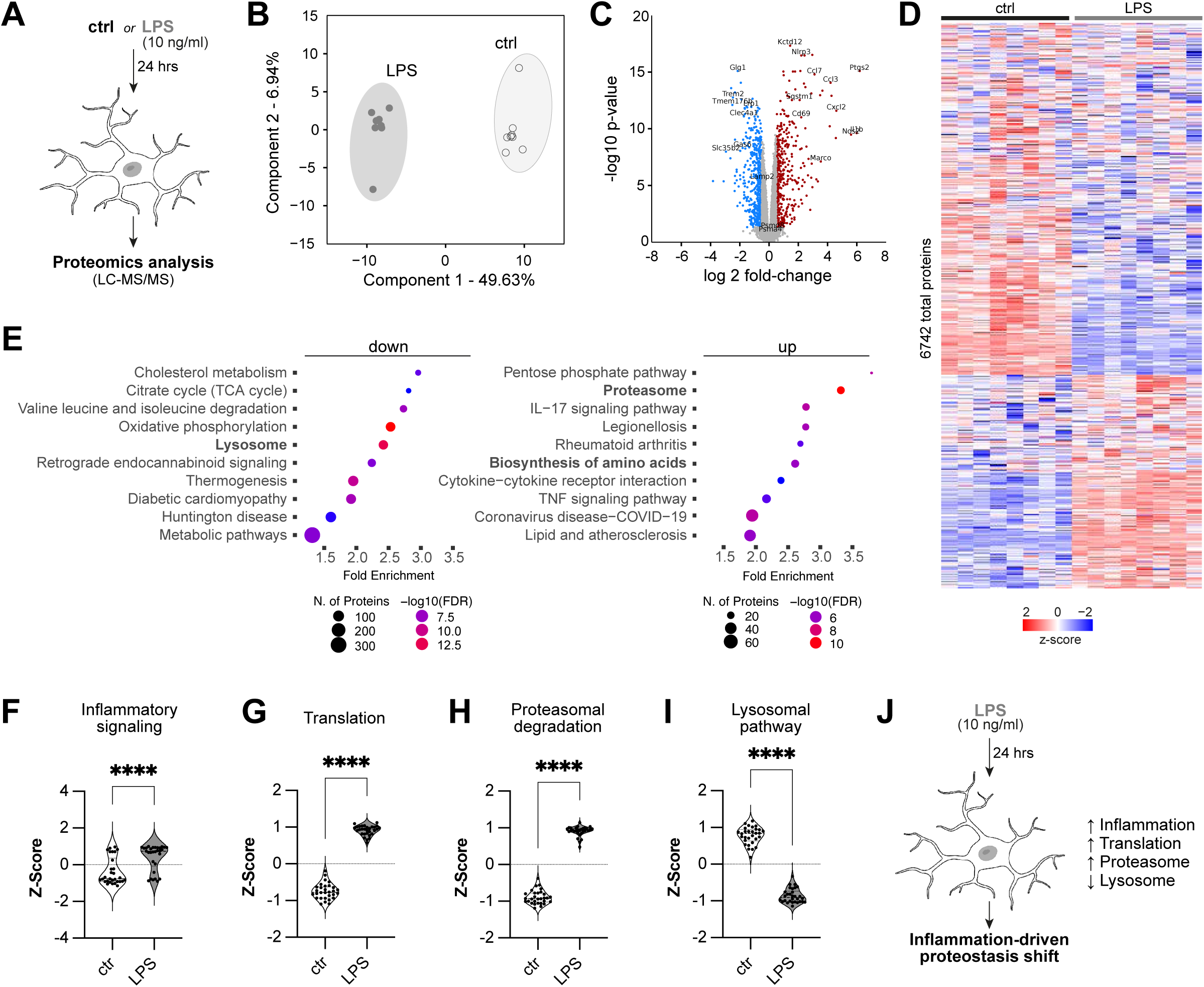
Inflammatory activation induces a global proteostasis shift in microglia. (A) Schematic overview of the proteomics workflow. Primary mouse microglia were treated with LPS (10 ng/ml) or left untreated (ctrl) for 24 h prior to proteomic analysis. (B) Principal component analysis (PCA) of the global proteome comparing LPS-treated (10 ng/ml) and untreated primary microglia. n=8. (C) Volcano plot showing differential protein abundance between LPS-treated and untreated microglia. n=8. (D) Heatmap showing Z-score-normalized protein abundance of all 6,742 detected proteins across individual samples. (E) Lollipop plot showing the top 10 up- and downregulated KEGG pathways following LPS treatment compared to ctrl. (F-I) Module scores of four selected Gene Ontology (GO) pathways related to inflammatory signaling (F), translation (G), proteasomal degradation (H), and lysosomal function (I) in LPS-treated and untreated control microglia. p<.0001 = ****. (J) Schematic summary of proteomic changes following LPS stimulation of primary microglia. Schematics were created with BioRender.com and Adobe Illustrator. Data are presented as violin plots showing the distribution of individual data points, with median and interquartile range indicated and were analyzed by t-test (F, G, H, I).

### TLR4 activation impairs degradation of extracellular α-Syn PFFs without strongly affecting uptake

We therefore investigated whether inflammatory activation affects the ability of microglia to process extracellular α-Syn and Tau aggregates, whose accumulation is a hallmark of several neurodegenerative diseases. To this end, we generated α-Syn and Tau pre-formed fibrils (PFFs) using established protocols and confirmed fibril formation before use (Figure S1). For α-Syn (as well as tau) experiments, we used 1 µM PFFs, a concentration that did not measurably change the microglial proteome in the absence of LPS (Figure S2). Proteins linked to phagocytic uptake were largely unchanged after LPS treatment (Figure 2A). We therefore first looked at how α-Syn enters microglia. Fluorescently labelled α-Syn PFFs were efficiently taken up and showed strong co-localization with 70-kDa dextran, a marker of macropinocytosis (Figures 2B-C). Internalized α-Syn was also found in LysoTracker-positive acidic compartments, showing that it enters the endolysosomal pathway. Blocking macropinocytosis with 5-[N-ethyl-N-isopropyl]amiloride (EIPA) strongly reduced uptake of both α-Syn and dextran, whereas inhibition of actin-dependent phagocytosis with cytochalasin D had only a small effect (Figure S3A-B). This indicates that macropinocytosis is the main uptake route for α-Syn PFFs under our conditions.

**Figure 2.**
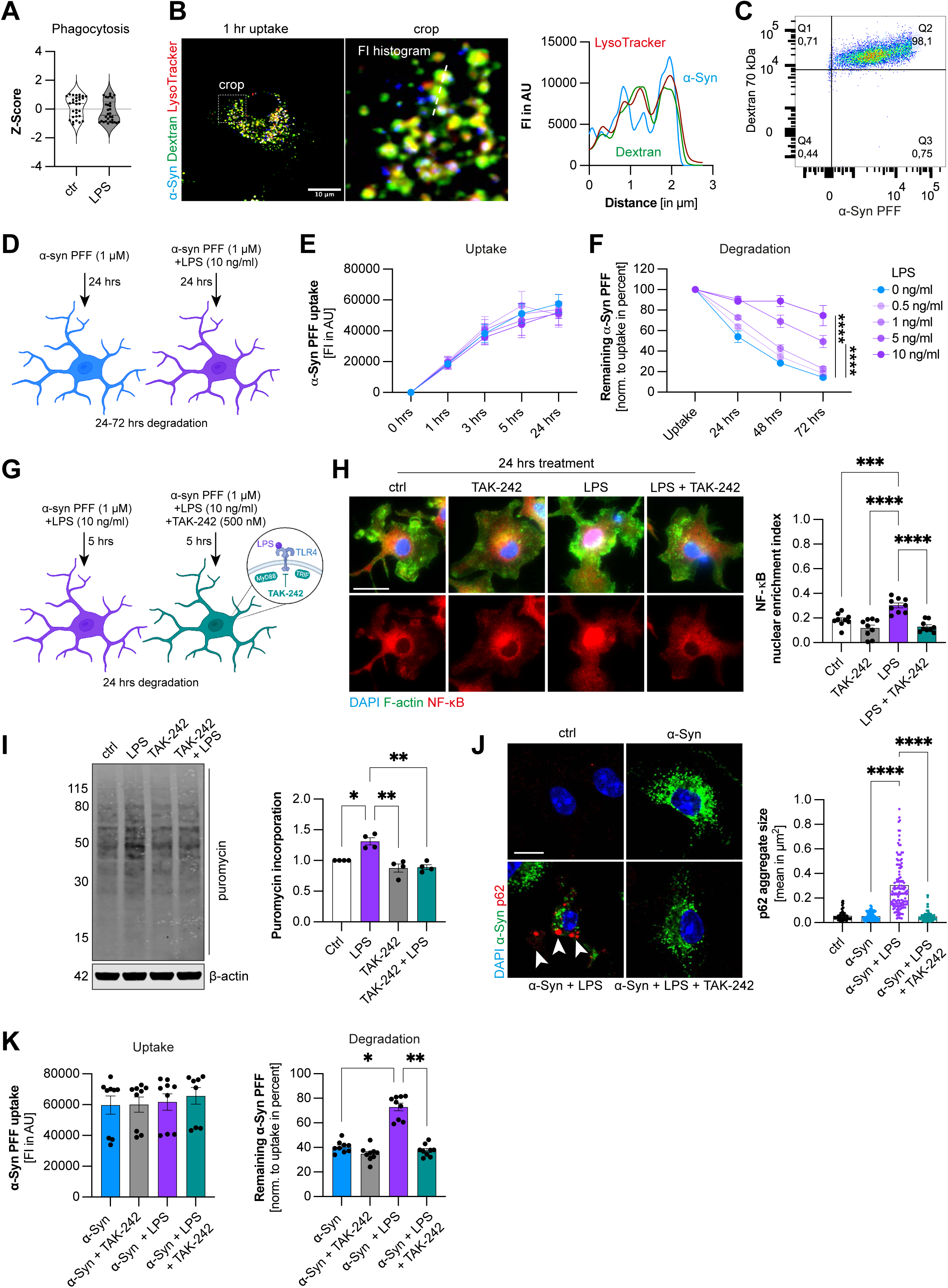
Inflammatory activation impairs microglial clearance of α-Syn PFFs. (A) Module scores of the Gene Ontology (GO) phagocytosis pathway in LPS-treated and untreated control microglia. (B) Representative immunofluorescence image (left panel) of primary microglia following 1 hr uptake of α-Syn PFFs (blue) and 70 kDa dextran (green), counterstained with LysoTracker Deep Red (red). Fluorescence intensity profiles (right panel) along the indicated line show overlap of all three signals. (C) Representative flow cytometry dot plot with marginal histograms showing 98.1% double-positive microglia for α-Syn PFFs and 70 kDa dextran. (D) Schematic overview of the α-Syn PFF uptake and degradation workflow for subsequent flow cytometry, immunofluorescence, and Western blot analyses. (E) Time-dependent uptake of α-Syn PFFs by primary microglia exposed to increasing LPS concentrations (0-10 ng/ml), as quantified by flow cytometry. n = 3, N = 6. (F) Time-dependent degradation of α-Syn PFFs by primary microglia exposed to increasing LPS concentrations (0-10 ng/ml), as quantified by flow cytometry. p<.0001 = ****. n = 3, N = 6. (G) Schematic overview of the experimental workflow to assess the effects of TLR4 inhibition with TAK-242 (500 nM) on α-Syn PFF clearance. (H) Representative immunofluorescence images (left panel) and quantification of NF-κB nuclear enrichment (right panel) in primary microglia following 24 hrs of the indicated treatments. Cells were stained for NF-κB (red), F-actin (green), and DAPI (blue). Ctrl and LPS conditions are identical to those shown in Figure 3B, as the data were obtained from the same experiment. n=3, N=9 fields of view. p<.0.001 = ***, p<.0001 = ****. Scale bar, 20 µm. (I) Representative Western blot (left panel) and quantification (right panel) showing newly synthesized proteins assessed by puromycin incorporation under the indicated treatment conditions. p<.05 = *, p<.01 = **. n=4. (J) Representative immunofluorescence images (left panel) and quantification of p62-positive aggregate size (right panel) in primary microglia following 5 hrs uptake of α-Syn PFFs (green). Cells were stained for p62 (red) and DAPI (blue). Arrowhead point to p62 aggregates. p<.0001 = ****. n = 3, N = 90-143 aggregates. Scale bar, 20 µm. (K) Quantification of uptake and degradation of α-Syn PFFs by microglia. Degradation is presented as percentage to uptake. Kruskal-Wallis with Dunn’s post-hoc test and one-way ANOVA was calculated and compared by Tuckey’s post-hoc for all conditions p<.05 = *, p<.01 = **. n=3, N=9. Schematics were created with BioRender.com and Adobe Illustrator. All graphs are presented as mean ± SEM and were analyzed by one-way ANOVA followed by Tukey’s test (E, F, H, I) or Kruskal-Wallis test followed by Dunn’s multiple-comparisons test (J, K).

Having established macropinocytosis as the main uptake route, we then separated the effects of LPS on α-Syn internalization from its subsequent degradation. Microglia were exposed to α-Syn PFFs for 24 h in the presence of increasing concentrations of LPS (0-10 ng/ml), followed by removal of extracellular α-Syn and a chase in α-Syn- and LPS-free medium for 24-72 h (Figure 2D). LPS caused only a modest reduction in α-Syn uptake (Figures 2E). In contrast, intracellular α-Syn persisted during the chase period in LPS-treated microglia in a concentration-dependent manner (Figures 2F and S3C). Thus, inflammatory activation has relatively little effect on α-Syn internalization while it strongly impairs the subsequent clearance of already internalized aggregates. To confirm that the observed decrease in intracellular α-Syn reflected degradation defects rather than altered cell viability or aggregate secretion, we assessed cell death by SYTOX staining and LDH release and found no difference between control and LPS-treated cells (Figure S3D-E). Likewise, α-Syn levels in the cell culture supernatant remained unchanged throughout the chase period (Figure S3F), indicating that the reduced degradation capacity is not due to changes in aggregate secretion. Importantly, inflammatory activation (LPS, 10 ng/ml) similarly impaired degradation of Tau PFFs (Figure S3G), suggesting that the observed phenotype reflects a broader defect in microglial processing of extracellular protein aggregates rather than an α-Syn-specific mechanism.

Given the increased protein synthesis and impaired α-Syn degradation following LPS treatment, we asked whether inflammatory activation also promotes the sequestration of intracellular proteins. We therefore examined p62, a selective autophagy receptor involved in the sequestration of protein aggregates, and observed the formation of enlarged p62-positive bodies in LPS-treated microglia. Since p62-dependent sequestration of α-Syn has previously been reported^14^, we asked whether internalized α-Syn was recruited to these structures. However, α-Syn showed little to no co-localization with p62-positive bodies during either uptake or degradation (Figure S4A). Instead, newly synthesized proteins accumulated within these structures, as shown by puromycin labeling (Figure S4B-C). To exclude a direct role of p62 in α-Syn clearance, we reduced p62 expression by siRNA. Although p62 depletion prevented body formation, it did not affect α-Syn degradation (Figure S5A-D). Together, these findings suggest that p62 bodies are not a major destination for internalized α-Syn and are not required for its degradation, but rather reflect a parallel proteostatic response to inflammatory activation. We therefore used p62-body formation as a morphological readout of this response and asked whether both the proteostatic changes and the degradation defect depended on TLR4 signaling.

To address this, we inhibited TLR4 with TAK-242 (Figure 2G). LPS induced robust NF-κB nuclear translocation at both 30 min and 24 h post treatment, whereas TAK-242 prevented this response at both time points (Figure 2H and S5E). Consistent with the increased translation signature identified in our proteomic analysis, LPS treatment (24 h, 10 ng/ml) also increased nascent protein synthesis, while TAK-242 reduced it back to control levels (Figure 2I). In parallel, LPS induced the formation of enlarged p62-positive bodies, which was prevented by TLR4 inhibition (Figure 2J and S5F). Strikingly, TAK-242 restored α-Syn degradation to near-control levels in LPS-treated microglia (Figure 2K).

Together, these data identify TLR4 signaling as a key upstream driver of the degradation defect and show that blocking TLR4 not only reverses the associated translational and p62 response, but also restores α-Syn clearance.

### Increased intracellular proteostatic stress is sufficient to impair extracellular α-Syn degradation

Since TLR4 inhibition simultaneously normalized translation, prevented p62-body formation, and restored α-Syn degradation, we asked which part of this response was functionally responsible for the degradation defect. To separate increased protein synthesis from upstream inflammatory signalling, we used cycloheximide (CHX) to inhibit translation downstream of TLR4 activation (Figure 3A). While LPS-induced NF-κB nuclear translocation remained intact in the presence of CHX (Figure 3B and S5G), puromycin incorporation was strongly reduced, confirming effective inhibition of nascent protein synthesis (Figures 3C). This was accompanied by a marked reduction in p62-positive body formation (Figure 3D and S5H). Importantly, despite preserved NF-κB activation, CHX restored α-Syn degradation in LPS-treated microglia (Figure 3E). These findings indicate that increased protein synthesis downstream of TLR4 contributes critically to the impaired degradation phenotype.

**Figure 3.**
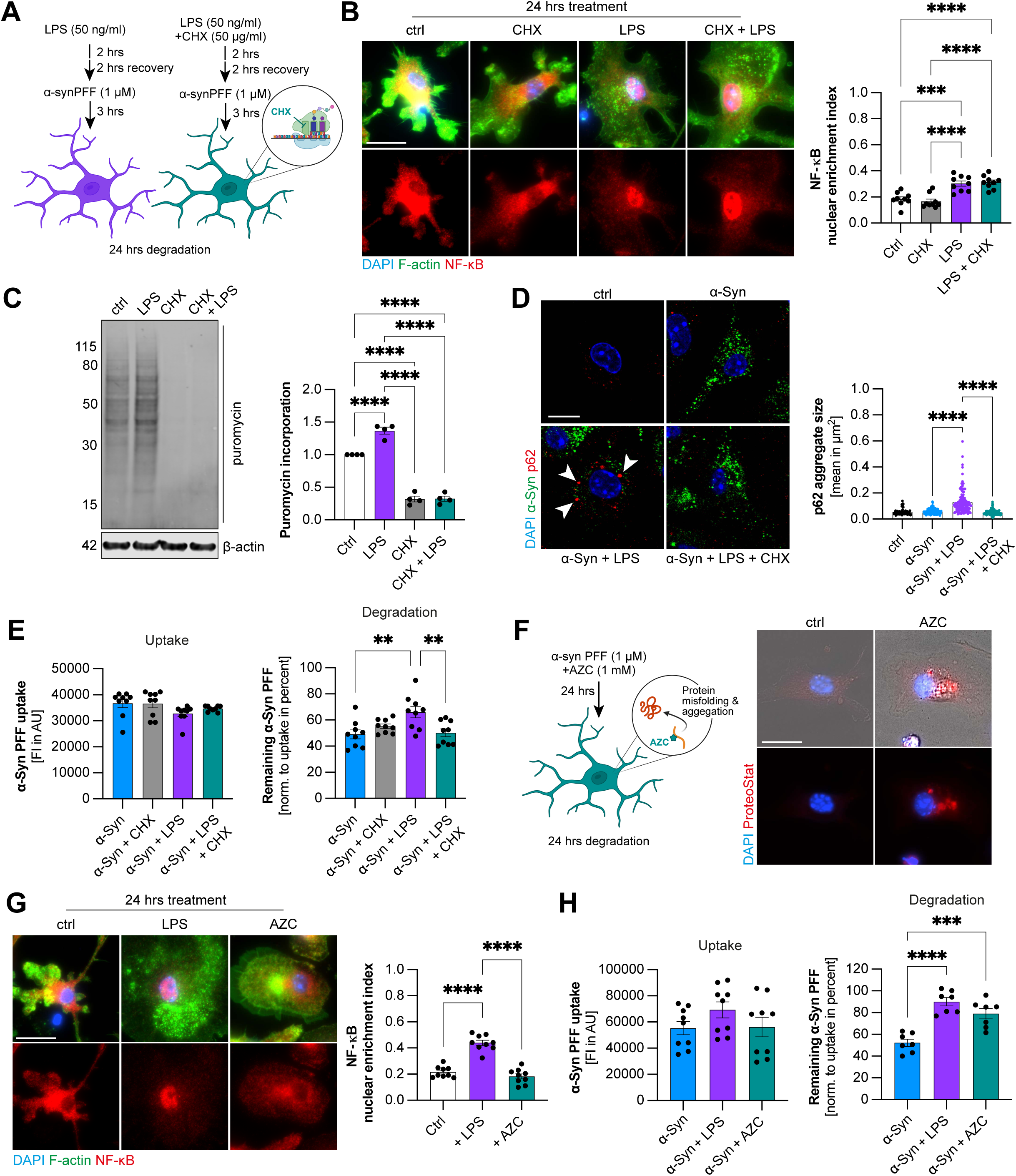
TLR4 activation promotes protein synthesis and impairs microglial α-Syn PFF clearance. (A) chematic overview of the experimental workflow to assess the effects of protein synthesis inhibition with cycloheximide (CHX, 50 µg/ml) on α-Syn PFF clearance. (B) Representative immunofluorescence images (left panel) and quantification of NF-κB nuclear enrichment (right panel) in primary microglia following 24 hrs treatment with CHX in the presence or absence of LPS. Cells were stained for NF-κB (red), F-actin (green), and DAPI (blue). Ctrl and LPS conditions are identical to those shown in Figure 2H, as the data were obtained from the same experiment. n=3, N=9 fields of view. p<.0.001 = ***, p<.0001 = ****. Scale bar, 20 µm. (C) Representative Western blot (left panel) and quantification (right panel) showing newly synthesized proteins assessed by puromycin incorporation under the indicated treatment conditions. p<.0001 = ****. n=4. (D) Representative immunofluorescence images (left panel) and quantification of p62-positive aggregate size (right panel) in primary microglia following the indicated treatments and 3 hrs uptake of α-Syn PFFs (green). Cells were stained for p62 (red) and DAPI (blue). Arrowheads point to p62 aggregates. p<.0001 = ****. n = 3, N = 90-143 aggregates. Scale bar, 20 µm. (E) Quantification of α-Syn PFF uptake (left panel) and degradation (right panel) in primary microglia under the indicated treatment conditions. Degradation was expressed as a percentage of the uptake measurement. Statistical significance was determined using one-way ANOVA followed by Tukey’s multiple-comparisons test. p<.01 = **. n=3, N=6. (F) Schematic overview (left panel) of the experimental workflow to assess the effects of proteostatic stress induced by L-azetidine-2-carboxylic acid (AZC, 1 mM), a proline analogue that promotes protein misfolding and aggregation. Representative image (right panel) of protein aggregation using ProteoStat dye (red) and DAPI (blue). Scale bar: 20 µm. (G) Representative immunofluorescence images (left panel) and quantification of NF-κB nuclear enrichment (right panel) in primary microglia following 24 h treatment with LPS or AZC. Cells were stained for NF-κB (red), F-actin (green), and DAPI (blue). n=3, N=9 fields of view. p<.0001 = ****. Scale bar, 20 µm. (H) Quantification of uptake (right panel) and degradation (left panel) of α-Syn PFFs by primary microglia. Degradation is presented as percentage to uptake p<.001 = ***, p<.0001 = ****. n=3, N=9. Schematics were created with BioRender.com and Adobe Illustrator. All graphs are presented as mean ± SEM and were analyzed by one-way ANOVA followed by Tukey’s test (B, E, G, H) or Kruskal-Wallis test followed by Dunn’s multiple-comparisons test (D).

The CHX experiments showed that reducing protein synthesis is sufficient to restore α-Syn degradation despite continued NF-κB activation. We therefore asked whether increased proteostatic stress itself is sufficient to induce the degradation defects. To test this, we used L-azetidine-2-carboxylic acid (AZC), a proline analogue that is incorporated into newly synthesized proteins and promotes protein misfolding (Figure 3F). In contrast to LPS, AZC did not induce NF-κB nuclear translocation at either 30 min or 24 h, indicating that the treatment did not trigger detectable NF-κB activation under these conditions (Figure 3G and S6A). Nevertheless, AZC impaired α-Syn degradation to a similar extent as LPS (Figure 3H). Thus, proteotoxic stress alone is sufficient to phenocopy the degradation defect without activating the NF-κB pathway. Together, these findings identify increased translational and proteostatic burden downstream of TLR4 as a key contributor to impaired α-Syn degradation and show that proteotoxic stress is sufficient to reproduce the phenotype independently of detectable NF-κB activation.

### Inflammatory activation preserves canonical autophagic flux while remodeling the endolysosomal pathway

Having established that increased translational and proteostatic burden contribute to impaired α-Syn degradation, we next asked how inflammatory activation affects the cellular degradative machinery (Figure 4A). In our proteomic analysis we found an upregulation of several autophagy-associated proteins following LPS stimulation, independent of α-Syn exposure (Figure S6B-C), while proteins associated with endosomal and lysosomal trafficking were reduced (Figure S6D).

**Figure 4.**
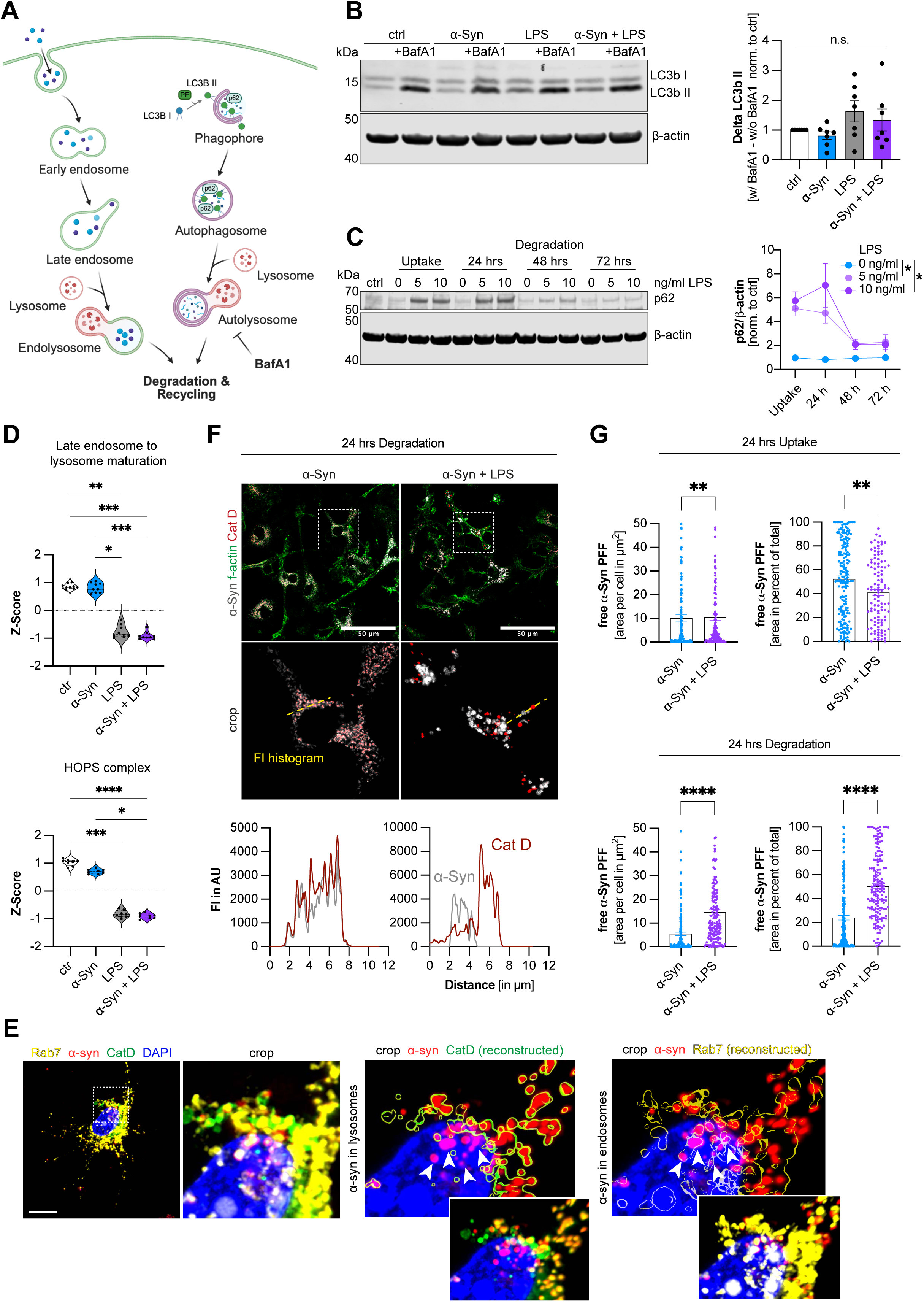
TLR4 activation preserves autophagic flux but impairs endolysosomal α-Syn processing. (A) Schematic overview of the canonical autophagy pathway and endolysosomal pathway. (B) Representative Western blot (left panel) and quantification (right panel) of autophagic flux assessed by LC3B-II accumulation in the presence or absence of bafilomycin A1 (BafA1, 100 nM, 4 hrs). LC3B-II protein levels were normalized to β-actin. n=7. (C) Representative Western blot (left panel) and quantification (right panel) of p62 protein levels in primary microglia following 24 hrs α-Syn PFF uptake and subsequent degradation for 24-72 h. p62 levels were normalized to β-actin. n=3. p<.05 = *. (D) Proteomics-derived module scores of the GO pathway “Late endosome to lysosome maturation” (upper panel) and “HOPS complex” (lower panel) under the indicated treatment conditions. p<.05 = *, p<.01 = **, p<.001 = ***, p<.0001 = ****. n=8. (E) Representative immunofluorescence images showing the subcellular localization of α-Syn PFFs (red), Rab7 (yellow), and cathepsin D (CatD, green) in primary microglia. Nuclei were counterstained with DAPI (blue). Three-dimensional reconstructions illustrate the spatial association of α-Syn PFFs with Rab7- and CatD-positive compartments, as indicated by arrowheads. Scale bar: 10 µm. (F) Representative immunofluorescence images (upper panel) of primary microglia following 24 hrs α-Syn PFF uptake (grey) in the presence or absence of LPS (10 ng/ml). Cells were stained for CatD (red) and F-actin (green). Fluorescence intensity profiles (lower panel) along the indicated lines show reduced colocalization of α-Syn PFFs with CatD-positive compartments following LPS treatment. Scale bars: 50 µm. (G) Quantification of α-Syn PFF puncta lacking CatD colocalization (free α-Syn puncta), expressed as absolute values and as a percentage of total α-Syn-positive area per cell. Measurements were performed following 24 hrs uptake (upper panel) and an additional 24 hrs degradation (lower panel) period under the indicated treatment conditions. p<.01 = **, p<.0001 = ****. n=3, N=104-199. Schematics were created with BioRender.com and Adobe Illustrator. All graphs are presented as mean ± SEM and were analyzed by t-test (G), one-way ANOVA followed by Tukey’s test (B, C) or Kruskal-Wallis test followed by Dunn’s multiple-comparisons test (D).

Hence, we first examined whether the increase in autophagy-associated proteins was accompanied by changes in the canonical autophagic flux. LC3B-II accumulation following 4 h treatment with the autophagy inhibitor bafilomycin A1 tended to be higher in LPS-treated microglia, although this difference was not statistically significant (Figure 4B and S6E). These data indicate that the autophagic flux remains largely preserved under inflammatory conditions. LPS also increased total p62 levels and induced p62-positive bodies. However, these structures declined during prolonged incubation together with total p62 abundance (Figure 4C and S6F), further indicating that intracellular protein turnover was not globally blocked under inflammatory conditions.

We next examined the endolysosomal pathway involved in the trafficking of α-Syn PFFs to the lysosomes. Proteomic analysis showed reduced abundance of proteins associated with late endosomal trafficking and lysosomal fusion, including several components of the HOPS complex (Figures 4D). Because HOPS also contributes to autophagosome-lysosome fusion, these changes cannot be considered specific to the endosomal pathway. Nevertheless, the overall proteomic pattern suggested a stronger reduction of proteins linked to endosome-lysosome trafficking than of the broader machinery supporting canonical autophagy (Figure S6G).

To relate these proteomic changes to α-Syn processing, we followed the intracellular localization of α-Syn PFFs along the late endosomal-lysosomal pathway. Internalized α-Syn was found in both Rab7-positive late endosomal compartments and CatD-positive lysosomes (Figure 4E). Notably, even after 24 h of degradation, some remaining α-Syn aggregates were still found in Rab7-positive compartments without detectable association with CatD. This suggests that a fraction of internalized α-Syn can persist in late endosomal compartments rather than progressing to CatD-positive degradative lysosomes. We therefore quantified the association of α-Syn with CatD-positive compartments. In α-Syn-treated microglia, the association of α-Syn with CatD-positive compartments increased during the degradation period (Figure 4F). In α-Syn+LPS-treated cells, this association was markedly reduced. Notably, LPS treatment also reduced total CatD abundance, suggesting that the decreased association may reflect reduced availability of CatD-positive degradative compartments, impaired trafficking of α-Syn to these compartments, or a combination of both. Consistent with this, the pool of α-Syn that was not associated with CatD-positive compartments declined during the subsequent degradation period in control microglia, whereas it remained largely unchanged in LPS-treated cells (Figure 4G). Together, these findings suggest that inflammatory activation limits the efficient delivery and/or processing of internalized α-Syn within CatD-positive degradative compartments, contributing to its impaired clearance.

Together, these findings show that inflammatory activation does not cause a general loss of canonical autophagic flux. Instead, it is associated with reduced abundance of endolysosomal and lysosomal trafficking machinery and reduced access of internalized α-Syn to CatD-positive degradative compartments.

### Pharmacological targeting of lysosomal regulation restores aggregate degradation during inflammation

Our proteomic analysis identified lysosomal proteins among the most significantly downregulated protein group following LPS-induced inflammatory activation (Figures 1 and 5A), suggesting that altered lysosomal protein expression may contribute to impaired extracellular α-Syn degradation. Consistent with this, western blot analysis confirmed decreased CatD abundance in LPS-treated microglia (Figure 5B), while proteomic analysis revealed reduced levels of TFEB, a master regulator of lysosomal biogenesis and gene expression (Figure 5C). We therefore asked whether pharmacologically targeting TFEB-dependent lysosomal regulation could alleviate the degradation defect under inflammatory conditions.

**Figure 5.**
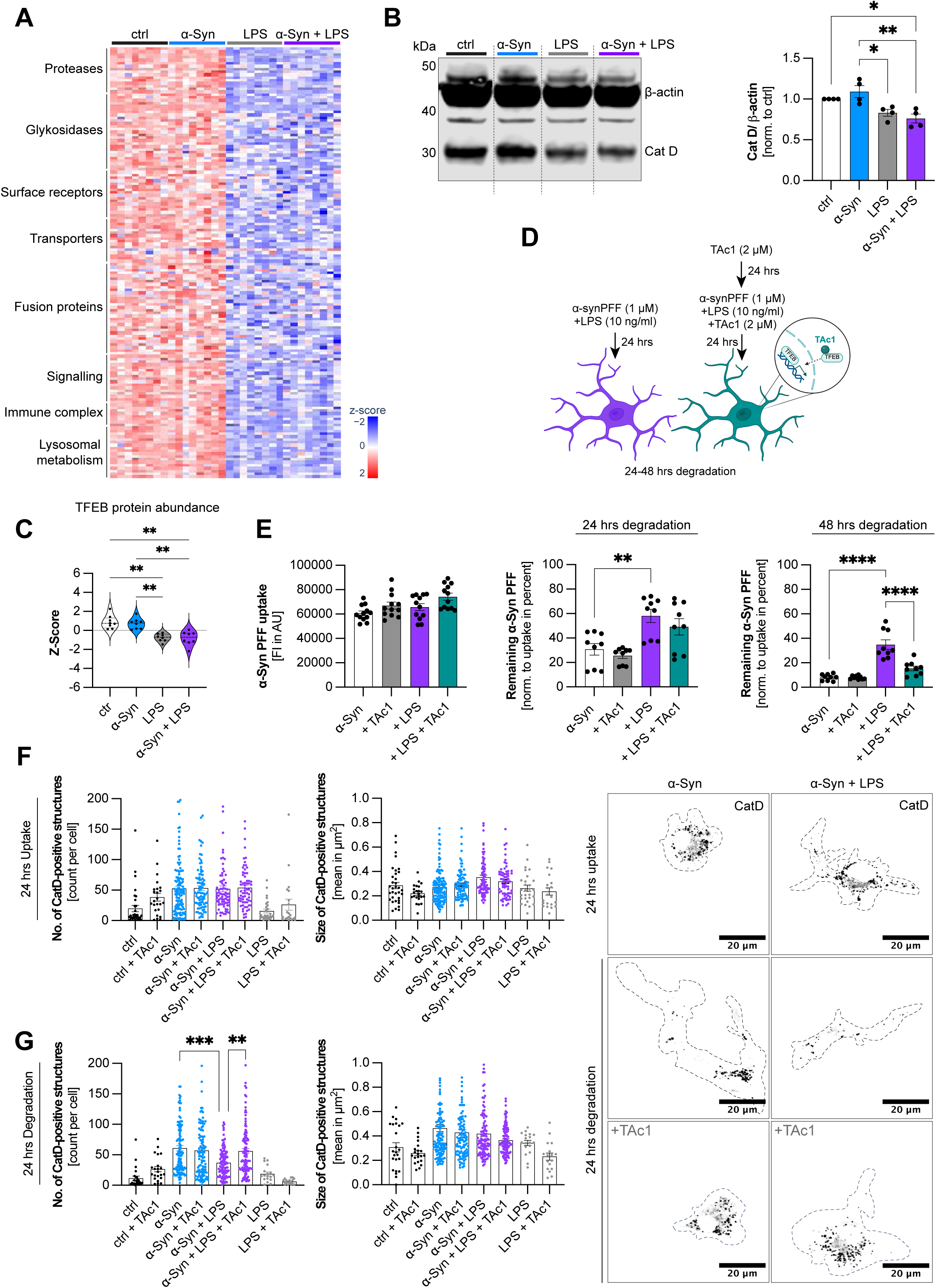
TFEB activation rescues inflammation-induced defects in α-Syn clearance. (A) Heatmap showing Z-score-normalized protein abundance of all proteins associated with the GO term “Lysosome” under the indicated treatment conditions. (B) Representative Western blot (left panel) and quantification (right panel) of CatD protein levels in primary microglia under the indicated treatment conditions. CatD levels were normalized to β-actin. Cropped regions of the Western blot are indicated by dashed lines. p<.05 = *, p<.01 = **. n=4. (C) Proteomics-derived TFEB protein abundance under the indicated treatment conditions, expressed as Z-scores. p<.01 = **. (D) Schematic overview of the experimental workflow to assess the effects of pharmacological TFEB activation on α-Syn PFF clearance. (E) Quantification of uptake (left panel) and degradation after 24 hrs (middle panel) and 48 hrs (right panel) of α-Syn PFFs by microglia. Degradation is presented as percentage to uptake. p<.01 = **, p<.0001 = ****. (F, G) Representative immunofluorescence images (right panel)) and quantification (left panel) of CatD-positive structures in primary microglia under the indicated treatment conditions. p<.01 = **, p<.001 = ***, p<.0001 = ****. n=3, N=34-118 (uptake) and N=17-125. Schematics were created with BioRender.com and Adobe Illustrator. All graphs are presented as mean ± SEM and were analyzed by one-way ANOVA followed by Tukey’s test (B, E (uptake data)) or Kruskal-Wallis test followed by Dunn’s multiple-comparisons test (C, E (degradation data), F, G).

To test this, microglia were treated with TFEB activator 1 (TAc1) during inflammatory stimulation (Figure 5D). TAc1 caused a modest reduction in LPS-induced NF-κB nuclear translocation after 30 min, but this effect was no longer detectable after 24 h, when NF-κB localization was comparable between LPS-and LPS+TAc1-treated cells (Figure S7A). TAc1 treatment restored α-Syn degradation in LPS-treated microglia following a 48 h chase period without significantly affecting α-Syn uptake (Figure 5E), suggesting that the rescue of a-Syn degradation primarily involves improved intracellular cargo processing rather than enhanced internalization. Similarly, TAc1 treatment restored the degradation of internalized Tau PFFs under inflammatory conditions (Figure S7B), indicating that the beneficial effect is not restricted to α-Syn. To further examine changes in the lysosomal compartment, we quantified CatD-positive structures during α-Syn exposure and the subsequent degradation period. Exposure to α-Syn PFFs increased the number of CatD-positive structures after 24 h, with no significant differences between treatment conditions at this time point (Figures 5F). However, following an additional 24 h degradation period, α-Syn-treated microglia retained a higher abundance of CatD-positive structures, whereas α-Syn+LPS-treated cells showed a reduction in both their number and the cellular area occupied by these structures (Figure 5G). Importantly, TAc1 treatment prevented this LPS-induced reduction, restoring both parameters to α-Syn-treated levels (Figure 5G). The delayed emergence of these differences suggests that inflammatory activation may compromise the maintenance of the CatD-positive compartment during prolonged aggregate processing rather than preventing its initial response to aggregate exposure. Together, these findings indicate that the inflammation-associated degradation defect is reversible and can be alleviated by pharmacological targeting of lysosomal regulatory pathways. Combined with the reduced abundance of lysosomal proteins, these observations support a model in which insufficient lysosomal adaptation contributes to impaired extracellular aggregate clearance during inflammatory activation.

### Features of impaired lysosomal α-Syn processing are present in human-relevant systems

To determine whether the proteostatic changes observed in primary mouse microglia are also seen in human cells, we reanalyzed two independent human datasets. In RNA-seq data from human brain-derived microglia stimulated with LPS (GSE249315)^18^, inflammatory pathways were strongly increased, whereas modules related to Rab7/late endosomes, the HOPS complex, SNARE-mediated fusion, lysosomal degradation, and TFEB/CLEAR signaling were reduced (Figure 6A). Autophagosome formation was also decreased in this dataset. Importantly, an independent proteomic dataset from iPSC-derived human microglia stimulated with LPS/IFN-γ showed the same overall pattern, with strong induction of inflammatory pathways accompanied by reduced lysosomal degradation, while the expression of autophagosome-associated proteins was not altered (Figure 6B)^19^. Together, these independent human datasets support the idea that inflammatory activation can be accompanied by a selective reduction in lysosomal programs rather than a uniform suppression of cellular degradation.

**Figure 6.**
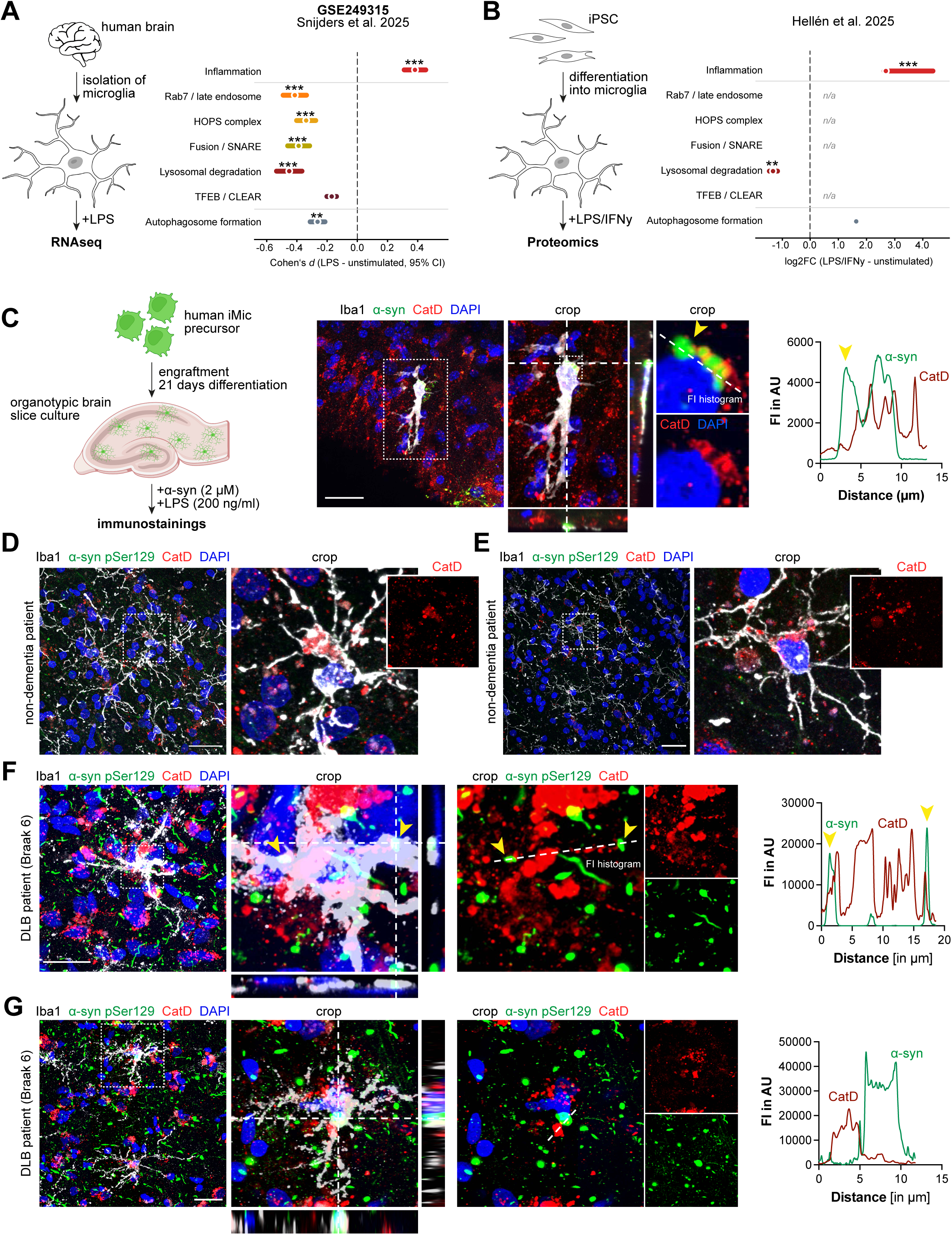
Inflammatory proteostasis alterations and α-Syn lysosomal localization in human microglia and brain tissue. (A) Reanalysis of publicly available RNA-seq data from human brain-derived microglia (GSE249315; Snijders et al., 2023) comparing LPS-stimulated and unstimulated cells. Transcriptional changes in the indicated biological processes are expressed as Cohen’s d (LPS vs. unstimulated) with 95% confidence intervals (CI). p<.01 = **, p<.001 = ***. (B) eanalysis of published proteomics data from human iPSC-derived microglia (Hellén et al., 2025) comparing LPS/IFNγ-stimulated and unstimulated cells. Changes in protein abundance are expressed as log2 fold change relative to unstimulated controls.p<.01 = **, p<.001 = ***. (C) Schematic overview (left panel) and representative immunofluorescence images (middel panel) of the engraftment of human iPSC-derived microglial precursors (iMic) into organotypic brain slice cultures. Cultures were treated with α-Syn (2 µM) and LPS (200 ng/ml) and stained for microglia (Iba1, white), α-Syn (green), CatD (red), and DAPI (blue). Fluorescence intensity profiles (left panel) illustrate the limited overlap of α-Syn and CatD signals. Scale bar, 30 µm. (D-E) Representative immunofluorescence images of post-mortem human brain tissue from neurologically healthy controls. Sections were stained for microglia (Iba1, white), phosphorylated α-Syn pSer129 (green), CatD (red), and DAPI (blue). Scale bar, 30 µm. (F-G) Representative immunofluorescence images (left panel) of post-mortem human brain tissue from patients with dementia with Lewy bodies (DLB, Braak stage 6). Sections were stained for Iba1 (white), α-Syn pSer129 (green), CatD (red), and DAPI (blue). Fluorescence intensity profiles (right panel) illustrate limited colocalization of pathological α-Syn with CatD-positive compartments. Scale bar, 30 µm. Schematics were created with BioRender.com and Adobe Illustrator. All graphs are presented as horizontal dot plot of module median Cohen’s d and log2 fold changes and were analyzed by paired t-test (A) and Fisher’s combined p-value (B).

We then examined α-Syn processing in a human-relevant tissue environment using chimeric slice cultures^61^. iPSC-derived human microglial precursors were engrafted into the slices and differentiated into mature microglia for 21 days. Slices were then exposed to α-Syn PFFs and LPS for 48 h. Internalized α-Syn could be readily detected within the engrafted human microglia, confirming uptake of extracellular aggregates in this tissue context. Notably, a fraction of α-Syn detected within microglia did not co-localized with CatD-positive compartments (Figure 6C), resembling the reduced α-Syn-CatD association observed in inflammatory primary microglia.

Finally, we analyzed post-mortem brain tissue from a non-demented control individual and a patient with dementia with Lewy bodies (DLB; Braak stage 6). Microglia in control tissue contained abundant CatD-positive lysosomal compartments, whereas microglia in DLB tissue contained prominent α-Syn deposits that showed little to no association with CatD-positive compartments (Figures 6D-G). While these static tissue images cannot directly measure lysosomal trafficking or degradation, they suggest that microglia-associated α-Syn can accumulate outside CatD-positive degradative compartments in human synucleinopathy.

Together, these findings show that key features of the phenotype identified in primary microglia extend to human systems: inflammatory activation is associated with reduced lysosomal programs, and microglia-associated α-Syn shows limited access to CatD-positive degradative compartments in human-relevant tissue models and DLB brain.

Collectively, our findings support a model in which inflammatory activation does not uniformly suppress microglial protein degradation (Figure 7). Instead, TLR4 signaling drives increased protein synthesis and an intracellular proteostatic demand while lysosomal adaptation becomes insufficient to maintain efficient processing of extracellular aggregate cargo. While the canonical autophagic flux remains largely preserved, aggregate-containing endolysosomal cargo shows reduced progression toward mature CatD-positive degradative compartments.

**Figure 7.**
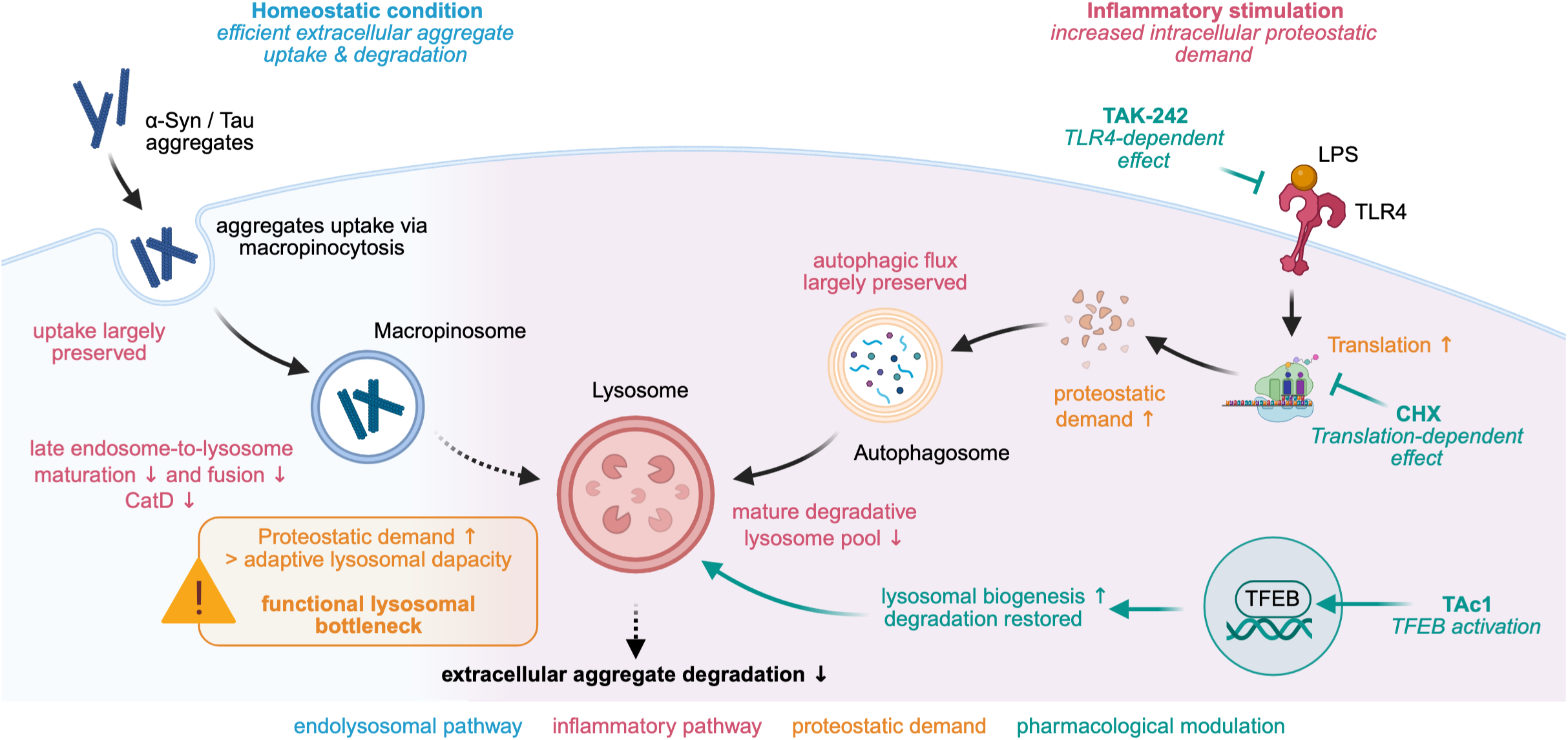
Working model of the inflammation-driven proteostasis shift in microglia. LPS-mediated activation of TLR4 increases protein synthesis and thereby elevates intracellular proteostatic burden in microglia. Although canonical autophagic flux remains largely preserved, lysosomal adaptation becomes insufficient to meet this increased demand. As a result, aggregate-containing endolysosomal cargo shows impaired progression toward mature CatD-positive degradative compartments, leading to reduced clearance of extracellular α-Syn aggregates. Proteostatic stress induced independently of TLR4 signaling is sufficient to phenocopy the α-Syn degradation defect, whereas increasing lysosomal capacity through TFEB activation restores aggregate clearance.

Independent induction of intracellular protein misfolding is sufficient to reproduce the α-Syn degradation defect, whereas increasing lysosomal capacity through TFEB activation restores aggregate clearance. Thus, inflammatory activation creates a demand-driven functional lysosomal bottleneck that uncouples intracellular proteostatic adaptation from efficient extracellular aggregate processing.

## Discussion

During neuroinflammation, microglia must simultaneously execute inflammatory effector programs while maintaining tissue homeostasis through the clearance of extracellular material, including misfolded protein aggregates^5,20^. Indeed, microglia can rapidly internalize and degrade α-Syn fibrils through endosomal and lysosomal compartments, underscoring the importance of efficient endolysosomal processing as a protective microglial function^8^. However, how these homeostatic clearance functions are coordinated with the proteostatic demands of inflammatory activation has remained poorly understood^21^. Here, we identify inflammatory proteostasis reprogramming as a mechanism that limits extracellular aggregate clearance. TLR4-dependent inflammatory activation impaired degradation, but not uptake, of extracellular α-Syn and Tau aggregates while simultaneously increasing protein synthesis and intracellular proteostatic stress. Reducing protein synthesis restored α-Syn degradation despite preserved NF-κB activation, whereas TLR4-independent induction of proteotoxic stress was sufficient to reproduce the degradation defect. Despite this increased burden, canonical autophagic flux remained largely preserved, whereas proteins involved in endolysosomal maturation, the association of α-Syn with CatD-positive compartments, and lysosomal abundance were reduced. Enhancing lysosomal biogenesis through TFEB activation restored aggregate degradation. Together, these findings support a model in which inflammatory activation creates a mismatch between intracellular proteostatic demand and adaptive lysosomal capacity, thereby limiting extracellular aggregate processing without causing a generalized collapse of intracellular degradation. Importantly, our findings challenge the assumption that inflammatory changes in microglial clearance primarily reflect altered phagocytic activity. Instead, they demonstrate that microglia can retain their ability to internalize pathological proteins while losing the capacity to efficiently degrade them. This distinction suggests that neuroinflammation may contribute to the accumulation of pathogenic proteins not necessarily by preventing their uptake, but by compromising their subsequent clearance.

A central implication of these findings is that intracellular protein quality control and extracellular cargo degradation, although initiated through distinct trafficking routes, are functionally coupled at the level of the lysosome. Autophagic and endocytic cargo ultimately converge at lysosomes, which act as a shared degradative endpoint for intracellular and extracellular material^22,23^. An increase in intracellular proteostatic demand could therefore place additional pressure on this shared degradative system, particularly if lysosomal adaptation does not increase accordingly. Consistent with this model, inflammatory activation profoundly reorganized the microglial proteome. TLR4 activation coordinately induced inflammatory signaling, protein synthesis, and stress-response pathways, while proteins involved in lysosomal maintenance and endolysosomal maturation were reduced (Figure 1 and 4). This is consistent with the well-established role of TLR signaling in rapidly reprogramming cellular metabolism to support immune effector functions, including the synthesis of cytokines, chemokines, and antimicrobial proteins^24^. Our findings extend this framework by showing that this biosynthetic reprogramming is accompanied by reduced lysosomal and endolysosomal programs, allowing different degradative functions to become unequally affected during inflammatory stress.

Several findings point to increased translational and proteostatic burden as an important component of the degradation defect. TLR4 inhibition normalized the LPS-induced increase in protein synthesis and restored α-Syn degradation (Figure 2G-K). More directly, inhibition of translation with CHX rescued α-Syn degradation even though LPS-induced NF-κB nuclear translocation remained intact (Figure 3A-E). Conversely, AZC-induced protein misfolding impaired α-Syn degradation without detectable NF-κB activation (Figure 3F-H). Together, these complementary experiments indicate that increased intracellular proteostatic burden is sufficient to constrain extracellular aggregate degradation without detectable NF-κB activation.

In parallel, inflammatory activation induced p62-positive condensates closely resembling previously described aggresome-like induced structures (ALIS), which sequester newly synthesized, ubiquitinated, or misfolded proteins during proteotoxic stress^25–30^. Newly synthesized proteins accumulated within these structures in our experiments (Figure S4B-C), whereas internalized extracellular α-Syn showed little to no association with them (Figure S4A). Moreover, p62 depletion prevented body formation without altering α-Syn degradation (Figure S5A-D). Thus, p62-body formation appears to reflect the accompanying proteostatic response rather than being a direct cause of impaired extracellular aggregate clearance.

Crucially, this increased intracellular proteostatic burden did not produce a global collapse of degradative function. Canonical autophagic flux remained largely preserved as assessed by LC3B-II turnover in the presence of bafilomycin A1 (Figure 4B and S6E), showing that canonical autophagic processing remained functional under inflammatory conditions. This observation is consistent with previous studies showing that TLR4 signaling can maintain or even stimulate autophagy in macrophages as part of the adaptive response to inflammatory stress^31,32^. In contrast, the endolysosomal machinery was more strongly altered. Proteomic analysis revealed reduced abundance of proteins involved in late endosome-to-lysosome maturation, including components of the HOPS tethering complex (Figure 4D). HOPS mediates tethering and fusion of late endosomes with lysosomes by bridging late-endosomal and lysosomal GTPase machinery^33^. HOPS also participates in autophagosome-to-lysosome fusion through interaction with the autophagosomal SNARE syntaxin 17 (STX17)^34^, indicating that individual fusion components are not exclusively pathway-specific. Nevertheless, our proteome-wide comparison revealed a preferential reduction of machinery associated with endosome-to-lysosome maturation and fusion, whereas the broader machinery supporting autophagosome-to-lysosome fusion was comparatively less affected (Figure S6G).

Consistent with these proteomic changes, inflammatory activation reduced the association of internalized α-Syn with CatD-positive compartments. Because LPS treatment also reduced total CatD abundance, the decreased association of α-Syn with CatD-positive compartments may reflect altered trafficking, reduced CatD availability, or a combination of both. Nevertheless, the combined proteomic and imaging data are consistent with less efficient progression of α-Syn-containing compartments toward mature degradative lysosomes. This is particularly relevant for α-Syn processing, as CatD has been identified as a major lysosomal protease mediating α-Syn degradation^35^. Together, these findings suggest that extracellular aggregate processing is more vulnerable to inflammatory proteostasis reprogramming than canonical autophagic flux.

These findings do not directly demonstrate competition between intracellular and extracellular cargo for individual lysosomes. Rather, they support a model in which inflammatory activation differentially affects the pathways converging on the lysosomal system, leaving canonical autophagic processing comparatively preserved while rendering endolysosomal processing of extracellular cargo particularly vulnerable when lysosomal adaptation is insufficient. A similar functional uncoupling has been described in microglia during ischemic stress, where increased autophagic activity can coexist with impaired engulfment and degradation of neuronal debris^36^. Although the upstream stressors differ, these observations support the broader idea that intracellular stress adaptation and extracellular clearance can become uncoupled under disease-associated stress.

The dynamics of the lysosomal compartment further support this model. Exposure to α-Syn initially expanded the pool of CatD-positive lysosomes irrespective of inflammatory stimulation, indicating that microglia can mount an early adaptive lysosomal response to increased extracellular cargo (Figure 5F). During the subsequent degradation period, however, α-Syn-treated microglia maintained this expanded lysosomal compartment, whereas α-Syn+LPS-treated cells failed to sustain it (Figure 5G). Thus, inflammatory microglia can initiate a lysosomal response to aggregate uptake but fail to maintain it over time. Our observation of reduced TFEB abundance following LPS stimulation (Figure 5C) is consistent with recent evidence identifying an IKK-β-TrCP2-dependent pathway controlling TFEB stability. Xiong et al. demonstrated that LPS promotes IKK-dependent degradation of TFEB and that prolonged LPS exposure reduces lysosomal compartments in an IKK-dependent manner in mouse embryonic fibroblasts, HeLa and HEK293T cells^37^. This provides a potential mechanism linking inflammatory signaling to reduced lysosomal adaptation, although whether the same pathway operates in microglia remains to be established. Consistent with this interpretation, treatment with the putative TFEB activator TAc1 restored the abundance of CatD-positive structures and rescued degradation of both α-Syn and Tau PFFs without altering phagocytic activity (Figures 5D-G and S7B). The rescue of both aggregates suggests that the limiting step is not α-Syn-specific. Our findings therefore extend the established role of TFEB in lysosomal biogenesis and proteostatic clearance to extracellularly acquired pathogenic aggregates^38–44^.

The relevance of this proteostatic shift was further supported across several human systems. Reanalysis of RNA-seq data from human brain-derived microglia showed strong induction of inflammatory pathways following LPS stimulation, together with reduced Rab7/late-endosomal, HOPS, SNARE/fusion, and lysosomal degradation signatures, while TFEB/CLEAR signatures also tended to decrease (Figure 6A)^18^. An independent proteomic dataset from iPSC-derived human microglia exposed to LPS/IFNγ showed a similar pattern, with strong inflammatory activation and reduced lysosomal degradation, whereas autophagosome-associated proteins were not reduced (Figure 6B)^19^. Although these datasets differ in experimental design, both support the idea that inflammatory activation can be accompanied by a disproportionate reduction in lysosomal programs rather than uniform suppression of degradative pathways.

We further observed α-Syn uptake by iPSC-derived human microglia engrafted into chimeric slice cultures after exposure to α-Syn PFFs and LPS (Figure 6C). A fraction of the α-Syn detected within the engrafted human microglia was not associated with CatD-positive compartments, showing that extracellular aggregate cargo can remain outside CatD-positive lysosomes in a human microglial tissue context. Finally, microglial α-Syn deposits in human DLB tissue similarly showed little association with CatD-positive compartments (Figure 6D-G). These observations cannot establish impaired lysosomal flux or causality, particularly given the static nature and limited scope of the post-mortem analysis, but provide human disease-relevant support for the phenotype identified experimentally.

The present findings are particularly relevant in the context of Parkinson’s disease (PD), where chronic neuroinflammation and impaired proteostasis are increasingly recognized as converging pathological processes^1,11^. Activated microglia and enhanced TLR4 signaling are consistently observed in the Parkinsonian brain and experimental disease models, where they precede or accompany progressive α-Syn pathology^11,14,45–53^. Our findings provide a potential mechanistic link between these processes by showing that inflammatory activation can preserve aggregate uptake while impairing subsequent degradation. The importance of microglial aggregate handling is further supported by recent *in vivo* work showing that engrafted human microglia restrict α-Syn aggregation and propagation in a PFF-based mouse model^54^. The use of LPS as an inflammatory stimulus may also have disease relevance beyond its role as a defined TLR4 agonist. The endotoxin hypothesis of PD proposes that gut dysbiosis and increased intestinal permeability may enhance systemic exposure to bacterial LPS in a subset of patients, thereby promoting peripheral and central inflammation^9^. Importantly, endotoxin exposure is unlikely to represent a sufficient or universal driver of PD, but may instead constitute an additional inflammatory hit that interacts with α-Syn pathology or other disease vulnerabilities.

Whether α-Syn fibrils directly activate TLR4 remains controversial. While several studies have reported TLR4-dependent inflammatory responses to α-Syn preparations^14,49^, others - including previous work from our laboratory - suggest that soluble monomeric and oligomeric α-Syn species are substantially more potent inflammatory stimuli than mature fibrils, acting predominantly through TLR2 and TLR5^55^. Consistent with this distinction, the sub-inflammatory concentrations of α-Syn PFFs used here did not elicit robust TLR4 activation (Figure S2). Recent work showed that α-Syn PFFs can directly impair lysosomal acidification and clearance in microglia, with TFEB activation restoring lysosomal acidification and α-Syn degradation^56^. In our model, by contrast, impaired α-Syn degradation emerged primarily during inflammatory activation and was associated with increased proteostatic demand, reduced abundance of endolysosomal maturation machinery, reduced association of α-Syn with CatD-positive compartments, and insufficient maintenance of lysosomal capacity despite largely preserved canonical autophagic flux. Our data therefore do not support a simple linear model in which fibrillar α-Syn activates TLR4 and thereby inhibits its own degradation. Rather, they suggest that inflammatory activation and extracellular aggregate accumulation can arise as parallel pathological processes that converge at the level of microglial proteostasis. In this setting, a pre-existing inflammatory state may reduce microglial capacity to process extracellular aggregates once they are encountered.

This concept also links to the strong genetic association between PD and lysosomal dysfunction^57^. Multiple disease-associated genes, including LRRK2 and GBA1, affect lysosomal trafficking or proteostasis, suggesting that genetic vulnerability and inflammatory reprogramming may converge on insufficient lysosomal capacity.

Overall, our findings highlight the importance of distinguishing phagocytic uptake from degradative capacity when evaluating microglial function during neuroinflammation. Rather than broadly suppressing microglial clearance, inflammatory activation creates an imbalance between intracellular proteostatic demand and lysosomal capacity, allowing microglia to internalize pathological proteins while compromising their subsequent degradation. In the context of neurodegeneration, this may contribute to the persistence of pathogenic protein aggregates despite ongoing microglial engagement. Importantly, the ability of TFEB activation to restore aggregate degradation demonstrates that this impairment is reversible and suggests that strengthening lysosomal capacity, rather than simply enhancing phagocytic uptake or broadly suppressing inflammation, may represent a promising strategy to preserve microglial clearance in chronically inflamed tissue.

## Conclusion

In summary, our study identifies a demand-driven lysosomal bottleneck as a mechanism by which inflammation limits extracellular aggregate clearance. Inflammatory activation does not globally disable lysosomal degradation but creates a mismatch between intracellular proteostatic demand and adaptive lysosomal capacity. Importantly, our findings demonstrate that microglia can retain their ability to internalize pathological proteins while losing the capacity to efficiently degrade them. By linking inflammatory reprogramming to impaired α-Syn and Tau degradation, our study highlights lysosomal adaptation as a critical determinant of microglial clearance and suggests that restoring degradative capacity may represent a promising strategy to counteract protein accumulation in neurodegenerative diseases.

## Limitations of the study

Our study has several limitations. First, although LPS-mediated TLR4 activation provides a well-defined and experimentally tractable model to dissect inflammatory signaling, it does not fully recapitulate the chronic and multifactorial inflammatory environment present in PD. Future studies using disease-relevant models will therefore be important to determine the generalizability of the proposed mechanism. Second, while our data implicate TLR4-dependent translational reprogramming as a critical driver of the lysosomal bottleneck, the signalling pathways linking inflammatory activation to increased proteostatic demand and impaired lysosomal adaptation remain incompletely understood. Defining these regulatory mechanisms will be important for understanding how inflammatory programs reshape microglial proteostasis. Third, our model proposes that increasing intracellular demand functionally limits extracellular aggregate degradation by exceeding the adaptive capacity of the lysosomal system. Although this interpretation is supported by the restoration of α-Syn degradation following TFEB activation, the precise mechanisms by which intracellular and extracellular cargo may compete for lysosomal resources remain to be directly resolved. Furthermore, although our findings reveal a functional uncoupling between aggregate uptake and degradation, the specific trafficking or degradative steps responsible for this impairment remain to be determined. Future studies combining quantitative analyses of lysosomal trafficking, cargo flux, and organelle dynamics will be important to further define these relationships. Finally, although our observations are supported by analyses of human postmortem tissue demonstrating altered spatial relationships between α-Syn aggregates and lysosomal markers, additional validation in larger human cohorts and disease-relevant experimental models will be necessary to establish the physiological relevance of the proposed mechanism and determine whether impaired lysosomal adaptation represents a general feature of neurodegenerative disease.

## Resource Availability

### Lead Contact

All data generated and/or analyzed during this study are either included in this article (and its Supplementary Information) or are available from the corresponding author at reasonable request. Further information and requests for resources and reagents should be directed to and will be fulfilled by the corresponding author, Hannah Scheiblich.

### Material Availability

This study did not generate new unique reagents.

### Data and Code Availability

The mass spectrometry proteomics data have been deposited to the ProteomeXchange Consortium via the PRIDE^58^ partner repository with the dataset identifier PXD085675.

## Acknowledgements

This work was supported by fundings from the Alzheimer Forschung Initiative (#23002R), the Gerhard und Ilse Schick Stiftung, the Deutsche Forschungsgemeinschaft (DFG, German Research Foundation) - 548919474 and the Fritz Thyssen Stiftung (Az. 10.25.1.027MN) to H.S. The authors would like to thank Sophie Schönen for her help regarding image analysis. We are deeply grateful to the patients and their families for their generous tissue donation. Biomaterial was provided by the DZNE Brain Bank of the German Center for Neurodegenerative Diseases (DZNE). We thank the FACS & Imaging Facility of the Max Planck Institute for Biology of Ageing (RRID:SCR_021354) for access to flow cytometry and microscopy infrastructure, as well as for technical support, image analysis resources, and data analysis software. We are grateful to the Proteomics Core Facility of the Max Planck Institute for Biology of Ageing for expert advice, mass spectrometry measurements, and proteomic data analysis. We also thank the Animal Facility of the Max Planck Institute for Biology of Ageing, in particular Laura Gobbers, for excellent animal husbandry. Finally, we acknowledge the Imaging Facility at CECAD, in particular Katrin Seidel and Felix Gaedke, for their expert assistance with electron microscopy. Schematics were created using BioRender (Scheiblich, H. (2027) https://BioRender.com/38aexa7) and Adobe Illustrator.

## Author contributions

Conceptualization: F.E. and H.S., Methodology: F.E. and H.S., Validation: F.E., L.W. and H.S., Formal analysis: F.E. and H.S., Investigation: F.E., L.W., A.He., L.E. and H.S., Resources: A.Ha. and H.S., Writing - original draft: F.E. and H.S., Writing - review & editing: H.S., Visualization: F.E. and H.S., Supervision: H.S., Funding acquisition: H.S.

## Declaration of generative AI and AI-assisted technologies in the writing process

During the preparation of this work, the authors used DeepL Write in order to improve readability and language as well as Heureka Bench in order to analyse proteomic datasets. After using these tools, the authors reviewed and edited the content as needed and take full responsibility for the content of the published article.

## Figure titles and legends

**Supplementary Figure 1 - Generation of pre-formed fibrils**

(A) Schematic representation of α-Syn pre-formed fibrils (PFFs) generation.

(B) Quantification of α-Syn fibrillization by Thioflavin T assay over 72 hrs. Raw (left panel) and normalized (right panel) data are presented.

(C) Representative western blot of sedimentation assay. α-Syn fibrils were sedimented by ultracentrifugation (100.000 xg) for 60 min. Low α-Syn signal in the supernatant indicates that the majority of α-Syn was present in an aggregated, sedimentable form.

(D) Electron microscopy image of α-Syn fibrils before and after sonication.

(E) Quantification of α-Syn PPFs length and thickness by electron microscopy.

(F) Schematic representation of Tau PFFs generation.

(G) Quantification of Tau fibrillization by Thioflavin T assay over 72 hrs. Raw and normalized data are presented.

(H) Representative western-blot of sedimentation assay. Tau fibrils were sedimented by ultracentrifugation (100.000 xg) for 60 min. Low Tau signal in the supernatant indicates that the majority of Tau was present in an aggregated, sedimentable form.

(I) Electron microscopy image of Tau fibrils before and after sonication.

(J) Quantification of Tau length and thickness by electron microscopy.

(K) Quantification of α-Syn and Tau protein endotoxin levels by LAL assay. Schematics were created with BioRender.com and Adobe Illustrator. Related to Figure 2.

**Supplementary Figure 2 - α-Syn PFFs at 1 µM do not change the microglial proteome in the absence of LPS**

(A) Schematic overview of the proteomics workflow. Primary mouse microglia were treated with α-Syn preformed fibrils (PFFs) or left untreated for 24 h prior to proteomic analysis.

(B) Principal component analysis (PCA) of proteomic data from eight independent experiments comparing α-Syn PFF-treated and untreated primary microglia.

(C) Volcano plot showing differential protein abundance between α-Syn PFF-treated and untreated control microglia.

Schematics were created with BioRender.com and Adobe Illustrator. Related to Figure 1 and 2.

**Supplementary Figure 3 - Characterization of α-Syn uptake and degradation in primary microglia.**

(A) Representative flow cytometry gating strategy for quantification of intracellular α- Syn PFFs in primary microglia following uptake. Cells were gated based on size and granularity, followed by exclusion of doublets and dead cells.

(B) Representative fluorescence images (left panel) and quantification (right panel) of primary microglia following 1 h uptake of α-Syn PFFs in the presence or absence of cytochalasin D or EIPA, inhibitors of actin-dependent phagocytosis and macropinocytosis, respectively. p<.0001 = ****. n=3, N=6.

(C) Representative Western blot (upper panel) and quantification (lower panel) of intracellular α-Syn following 24 h uptake and subsequent degradation for 24-72 h. α-Syn signals were normalized to β-actin and expressed as a percentage of the signal measured immediately after uptake. p<.05 = *. n=3.

(D) Quantification of microglial viability based on SYTOX dead-cell staining measured by flow cytometry. n=3.

(E) Quantification of microglial viability based on lactate dehydrogenase (LDH) release into the culture supernatant following α-Syn PFF uptake and degradation. p<.0001 = ****. n=3.

(F) Representative Western blot (upper panel) and quantification (lower panel) of extracellular α-Syn in culture supernatants during the degradation period, as determined by Western blotting. α-Syn levels are presented as raw fluorescence intensity (FI).

(G) Flow cytometric quantification of intracellular tau PFFs in primary microglia following 24 h uptake in the presence or absence of LPS (10 ng/ml). Data were normalized to the uptake measurement. ****p< 0.0001. n=3, N=9.

All graphs are presented as mean ± SEM and were analyzed by t-test (G) or one-way ANOVA followed by Tukey’s test (B, C, D, E, F). Related to Figure 2.

**Supplementary Figure 4 - Internalized α-Syn does not localize to p62 bodies**

(A) Representative immunofluorescence images of primary microglia following 5 hrs uptake and 24 hrs degradation of α-Syn PFFs (green), counterstained for p62 (red) and DAPI (blue). Fluorescence intensity profiles of α-Syn and p62 along the indicated line show distinct signals with little overlap, indicating limited colocalization. Scale bars: 20 µm.

(B) Schematic illustration of puromycin pulse-labeling to visualize newly synthesized proteins and assess their localization within p62-positive condensates.

(C) Representative immunofluorescence images of primary microglia following 24 hrs exposure to α-Syn PFFs (green) with and without LPS (10 ng/ml) and subsequent treatment with puromycin (1 µM) for 1 hr before fixation. Cycloheximide (CHX, 100 µM) was applied 1 hr before puromycin treatment as a negative control for protein synthesis. Cells were stained for p62 (red), puromycin (white), and DAPI (blue). Magnified regions show colocalization of puromycin and p62 in α-Syn PFF-and LPS- treated microglia, indicating the accumulation of newly synthesized proteins within p62-positive condensates. Scale bars: 50 µm.

Schematics were created with BioRender.com and Adobe Illustrator.

**Supplementary Figure 5 - p62 depletion does not rescue α-Syn degradation**

(A) Schematic overview of siRNA-mediated knockdown workflow. Primary microglia were magnetofected with siRNA and treated 48 hrs later with with α-Syn PFFs (1 µM) and LPS (10 ng/ml) for 24 h.

(B) Representative Western blot (left panel) and quantification (right panel) showing p62 protein levels in untreated control (ctrl), scrambled siRNA (scramble), and p62 (SQSTM1) siRNA-transfected microglia. p62 protein levels were normalized to β-actin. One-way ANOVA and Tuckey’s post-hoc test p<.05 = *, p<.01 = **.

(C) Representative immunofluorescence images of primary microglia following 48 hrs of siRNA-mediated knockdown and subsequent treatment with α-Syn PFFs (1 µM) and LPS (10 ng/ml) for 5 h. Cells were stained for p62 (red) and DAPI (blue) to visualize p62-positive condensates. Scale bars: 50 µm.

(D) Flow cytometric quantification of α-Syn PFF uptake (left panel) and degradation (right panel) in primary microglia following 48 hrs of siRNA-mediated knockdown. Uptake was assessed after 24 hrs of α-Syn PFF exposure, while degradation was measured following an additional 24 hrs chase period. p<.05 = *. n=3, N=7.

(E) Representative immunofluorescence images (left panel) and quantification of NF-κB nuclear enrichment (right panel) in primary microglia treated with TAK-242 (500 nM) in the presence or absence of LPS (10 ng/ml) for 30 min. Cells were stained for NF-κB (red), F-actin (green), and DAPI (blue). Ctrl and LPS conditions are identical to those shown in Figure S5G and S7A, as the data were obtained from the same experiment. n=3, N=9 fields of view. p<.0001 = ****. Scale bar, 20 µm.

(F) Representative immunofluorescence images (left panel) and quantification of p62-positive puncta per cell (right panel) in primary microglia under control conditions (ctrl) or treated with α-Syn, α-Syn+LPS, or α-Syn+LPS+TAK-242. Cells were stained for α-Syn (green), p62 (red), and DAPI (blue). Crops are shown in Figure 3D. n=3, N=90-143. p<.0001 = ****. Scale bar, 50 µm.

(G) Representative immunofluorescence images (left panel) and quantification of NF-κB nuclear enrichment (right panel) in primary microglia treated with CHX (50 µg/ml) in the presence or absence of LPS (10 ng/ml) for 30 min. Cells were stained for NF-κB (red), F-actin (green), and DAPI (blue). Ctrl and LPS conditions are identical to those shown in Figure S5E and S7A, as the data were obtained from the same experiment. n=3, N=9 fields of view. p<.0001 = ****. Scale bar, 20 µm.

(H) Representative immunofluorescence images (left panel) and quantification of p62-positive puncta per cell (right panel) in primary microglia under control conditions (ctrl) or treated with α-Syn, α-Syn+LPS, or α-Syn+LPS+CHX. Cells were stained for α-Syn (green), p62 (red), and DAPI (blue). Crops are shown in Figure 3I. n=3, N=74-132. p<.0001 = ****. Scale bar, 50 µm.

All graphs are presented as mean ± SEM and were analyzed by one-way ANOVA followed by Tukey’s test (B, D, E, F, G, H). Schematics were created with BioRender.com and Adobe Illustrator. Related to Figure 2 and 3.

**Supplementary Figure 6 - LPS increases autophagy-associated proteins while impairing endolysosomal trafficking and fusion**

(A) Representative immunofluorescence images (left panel) and quantification of NF-κB nuclear enrichment (right panel) in primary microglia following treatment with AZC (1 mM) for 30 min. Cells were stained for NF-κB (red), F-actin (green), and DAPI (blue). n=3, N=9 fields of view. p<.0001 = ****. Scale bar, 20 µm.

(B) Z-score of the core autophagy module under the indicated treatment conditions (ctrl, α-Syn, LPS, α-Syn+LPS). p<.05 = *.

(C) Changes in protein abundance of the core autophagy machinery (ULK1 initiation complex, ATG12 and ATG8/LC3 conjugation systems, selective autophagy receptors, and membrane source/elongation machinery) following LPS treatment, expressed as log2 fold change relative to ctrl. Proteins are color-coded by functional category. p<.05 = *, p<.0.01 = **, p<.0.001 = ***.

(D) Changes in protein abundance of the PI3P/PIKfyve-dependent endolysosomal maturation pathway following LPS treatment, expressed as log2 fold change relative to ctrl. Proteins are color-coded by functional category. p<.05 = *.

(E) Western blot quantification of LC3B-I normalized to β-actin (left panel), LC3B-II normalized to β-actin (middle panel), and the LC3B-II/LC3B-I ratio (right panel) in primary microglia treated with α-Syn, LPS, or their combination in the presence or absence of bafilomycin A1 (Baf). p<.05 = *. n=7.

(F) Western blot quantification of p62 protein levels normalized to β-actin under the indicated treatment conditions (Ctrl, α-Syn PFFs, LPS, and α-Syn PFFs + LPS), in the presence or absence of Baf. Data were normalized to untreated controls. n=4.

(G) Changes in the abundance of proteins involved in autophagosome-lysosome fusion, endosome-lysosome fusion, and shared tethering machinery following LPS treatment, expressed as log2 fold change relative to untreated controls (LPS vs. Ctrl). Proteins are color-coded according to their functional categories. Statistical significance was determined using paired t-tests. p<.05 = *, p<.01 = **, p<.001 = ***, p<.0001 = ****.

All graphs are presented as mean ± SEM and were analyzed by moderated t-test (limma) (C, D, G) or one-way ANOVA followed by Tukey’s test (B, E). Related to Figure 3 and 4.

**Supplementary Figure 7 - TFEB activation restores Tau PFF degradation during inflammation**

(A) Representative immunofluorescence images (left panel) and quantification of NF-κB nuclear enrichment (right panel) in primary microglia following treatment with TAc1 (2 µM) in the presence or absence of LPS (10 ng/ml) for 30 min and 24 hrs. Cells were stained for NF-κB (red), F-actin (green), and DAPI (blue). Ctrl and LPS conditions are identical to those shown in Figure S4E+G (for 30 min) and Figure 2H and 3G (for 24 hrs), as the data were obtained from the same experiment. n=3, N=9 fields of view. p<.01 = **, p<.0001 = ****. Scale bar, 20 µm.

(B) Quantification of Tau PFFs in microglia after 24 hrs uptake (left panel) as well as 24 hrs (middle panel) or 48 hrs (right panel) degradation in the presence or absence of Tac1 (2 µM) as measured by flow cytometry. p<.05 = *, p<.01 = **, p<.0.001 = ***, p<.0001 = ****.

All graphs are presented as mean ± SEM and were analyzed by one-way ANOVA followed by Tukey’s test (A, B). Related to Figure 5.

## STAR Methods

### Key Resource Table

| **REAGENT or RESOURCE** | **SOURCE** | **IDENTIFIER** |
|---|---|---|
| **Antibodies** |  |  |
| Donkey anti-Goat IgG (H + L) Alexa Fluor 647 | Thermo Fisher Scientific | Cat#A32849TR; RRID: AB_2866498 |
| Donkey anti-Goat IgG (H+L) Alexa Fluor 488 | Thermo Fisher Scientific | Cat#A-11055; RRID: AB_2534102 |
| Donkey anti-Mouse IgG (H+L) Alexa Fluor 488 | Thermo Fisher Scientific | Cat#A-21202; RRID: AB_141607 |
| Donkey anti-Rabbit IgG (H+L) Alexa Fluor 594 | Thermo Fisher Scientific | Cat#A-21207; RRID: AB_141637 |
| Goat anti-Cathepsin D | Thermo Fisher Scientific | Cat#PA547046; RRID: AB_2609303 |
| Goat anti-Mouse IgG (H + L) Alexa Fluor 488 | Thermo Fisher Scientific | Cat#A28175; RRID: AB_2536161 |
| Goat anti-Mouse IgG (H + L) Alexa Fluor 594 | Thermo Fisher Scientific | Cat#A-11032; RRID: AB_2534091 |
| Goat anti-Rabbit IgG (H + L) Alexa Fluor 488 | Thermo Fisher Scientific | Cat#A-11008; RRID: AB_143165 |
| Goat anti-Rabbit IgG (H + L) Alexa Fluor 594 | Thermo Fisher Scientific | Cat#A-11012; RRID: AB_2534079 |
| Goat anti-Rabbit IgG (H + L) Alexa Fluor 647 | Thermo Fisher Scientific | Cat#A-21245; RRID: AB_2535813 |
| Goat anti-Rat IgG (H + L) Alexa Fluor 488 | Thermo Fisher Scientific | Cat#A-11006; RRID: AB_2534074 |
| IRDYE® 680LT GOAT ANTI-MOUSE IGG SECONDARY ANTIBODY | LI-COR Biosciences | Cat#926-68020; RRID: AB_10706161 |
| IRDYE® 680LT GOAT ANTI-RABBIT IGG SECONDARY ANTIBODY | LI-COR Biosciences | Cat#926-68021; RRID: AB_10706309 |
| IRDYE® 800CW GOAT ANTI-MOUSE IGG SECONDARY ANTIBODY | LI-COR Biosciences | Cat#926-32210; RRID: AB_621842 |
| IRDYE® 800CW GOAT ANTI-RABBIT IGG SECONDARY ANTIBODY | LI-COR Biosciences | Cat#926-32211; RRID: AB_621843 |
| Mouse anti-beta actin | Proteintech | Cat#66009-1-Ig; RRID: AB_2687938 |
| Mouse anti-Rab7 (E9O7E) | Cell Signaling Technology | Cat#95746; RRID: AB_2800252 |
| Mouse anti-STEM121 | Takara Bio | Cat#Y40410; RRID: AB_2801314 |
| Mouse anti-α-synuclein (clone 4B12/Synuclein) | BioLegend | Cat#807801; RRID: AB_2564730 |
| Mouse anti-α-synuclein phospho (Ser129) (clone P-syn/81A) | BioLegend | Cat#MMS-509; RRID: AB_2564891 |
| Rabbit anti-Iba1 | Wako | Cat#019–19741; RRID: AB_839504 |
| Rabbit anti-MAP1LC3b | Sigma-Aldrich | Cat#L7543-200UL; RRID: AB_796155 |
| Rabbit anti-NF-kappaB p65 (D14E12) | Cell Signaling Technology | Cat#8242; RRID: AB_10859369 |
| Rabbit anti-p62/SQSTM1 | Cell Signaling Technology | Cat#5114; RRID: AB_10624872 |
| Rabbit anti-puromycin | Abcam | Cat#ab315887; RRID: AB_3716527 |
| Ultra-LEAF™ Purified Rat anti-mouse CD115 (CSF-1R) | BioLegend | Cat#135539; RRID: AB_2832483 |
| **Bacterial and virus strains** |  |  |
| E.coli BL21 DE3 CodonPlus cells | Agilent Technologies | Cat#230245 |
| **Biological samples** |  |  |
| Human post-mortem brain tissue | Brain Bank DZNE | N/A |
| **Chemicals, peptides, and recombinant proteins** |  |  |
| 2-Mercaptoethanol (50 mM) | Thermo Fisher Scientific | Cat#31350010 |
| 2x MEM no phenolred | Thermo Fisher Scientific | Cat#21935028 |
| 4-(2-Hydroxyethyl)-1-piperazineethanesulfonic acid (HEPES) | Carl Roth | Cat#6763.1 |
| 5-(N-Ethyl-N-isopropyl)amiloride (EIPA) | Sigma-Aldrich | Cat#A3085 |
| Acetic acid | Supelco | Cat#5330010050 |
| Acetonitrile (ACN) | Supelco | Cat#1000291000 |
| Advanced DMEM/F12 | Thermo Fisher Scientific | Cat#12634010 |
| AggreWell™800, 24-well | Stemcell Technologies | Cat#34811 |
| Anti-Adherence Rinsing Solution | Stemcell Technologies | Cat#07010 |
| Ascorbic acid | Sigma-Aldrich | Cat#A4403-100mg |
| ATTO-405 NHS-ester | Thermo Fisher Scientific | Cat#A30000 |
| ATTO-488 NHS-ester | Atto-Tec GmbH | Cat#AD 488-35 |
| ATTO-647N NHS-ester | Sigma-Aldrich | Cat#18373 |
| Bafilomycin A1 | Sigma-Aldrich | Cat#B1793 |
| Bovine serum albumin (BSA) | Sigma-Aldrich | Cat#A8806 |
| BSA Fraction V, 7.5% Solution | Thermo Fisher Scientific | Cat#15260037 |
| Calcium chloride (CaCl2) | Sigma-Aldrich | Cat#21115-100ml |
| Chloroacetamide (CAA) | Thermo Fisher Scientific | Cat#A15238.30 |
| Cycloheximide | Carl Roth | Cat#8682.1 |
| D-Glucose | Carl Roth | Cat#X997.1 |
| Dextran, Fluorescein, 70,000 MW, Anionic, Lysine Fixable | Thermo Fisher Scientific | Cat#D1822 |
| DL-Dithiothreitol (DTT) | Sigma-Aldrich | Cat#D0632 |
| DNase 1 | Sigma-Aldrich | Cat#11284932001 |
| Dulbecco's Modified Eagle's Medium (DMEM) | Thermo Fisher Scientific | Cat#31966047 |
| Dulbecco's Phosphate-Buffered Saline (DPBS) | Thermo Fisher Scientific | Cat#14190169 |
| Formaldehyde | Science Services | Cat#E15700 |
| Formic acid 100% | Supelco | Cat#5330020050 |
| GlutaMAX | Thermo Fisher Scientific | Cat#3505-038 |
| Glutaraldehyde | Merck Millipore | Cat#G6257 |
| Guanidinium chloride | SAFC | Cat#1370371000 |
| HALT protease inhibitor | Thermo Fisher Scientific | Cat#78429 |
| Hank's Balanced Salt Solution (HBSS) | Thermo Fisher Scientific | Cat#14170120 |
| Heparin Sodium Salt | Sigma-Aldrich | Cat#H3149 |
| Horse serum, heat inactivated | Thermo Fisher Scientific | Cat#26050-088 |
| Human BMP-4, premium grade | Miltenyi Biotec | Cat#130-111-167 |
| Human IL-3, premium grade | Miltenyi Biotec | Cat#130-095-069 |
| Human M-CSF, premium grade | Miltenyi Biotec | Cat#130-096-493 |
| Human VEGF (165) IS, premium grade | Miltenyi Biotec | Cat#130-109-396 |
| Insulin (human recombinant) | Thermo Fisher Scientific | Cat#12585014 |
| L-azetidine-2-carboxylic acid (AZC) | MedChemExpress | Cat#HY-W050044 |
| Lipopolysaccharide from Escherichia coli K12 (LPS) | InvivoGen | Cat#tlrl-ek |
| Magnesium sulfate (MgSO4) | Sigma-Aldrich | Cat#83266 |
| Minimal Essential Medium (MEM) | Thermo Fisher Scientific | Cat#32360-26 |
| mTeSR™ Plus | Stemcell Technologies | Cat#100-0276 |
| N2-Supplement | Thermo Fisher Scientific | Cat#17502048 |
| Normal donkey serum | Jackson Immuno Research | Cat#017-000-121; RRID: AB_2337258 |
| Normal Goat Serum | Abcam | Cat#ab7481; RRID: AB_2716553 |
| NuPAGE™ MES SDS Running Buffer | Thermo Fisher Scientific | Cat#NP0002 |
| Opti-MEM™ I Reduced Serum Medium | Thermo Fisher Scientific | Cat#31985062 |
| Paraformaldehyde | Sigma-Aldrich | Cat#P6148 |
| Penicillin/Streptomycin | Thermo Fisher Scientific | Cat#15070063 |
| Phalloidin Alexa Fluor™ 647 | Thermo Fisher Scientific | Cat#A22287 |
| Phalloidin Alexa Fluor™ AlexaFluor488 | Cell Signaling Technology | Cat#8878 |
| Phalloidin Alexa Fluor™ TexasRed | Thermo Fisher Scientific | Cat#T7471 |
| PhosphoSTOP | Sigma-Aldrich | Cat#4906845001 |
| Poly-L-lysine Hybridomide | Sigma-Aldrich | Cat#P1524 |
| ProLong™ Diamond Antifade Mountant | Thermo Fisher Scientific | Cat#P36965 |
| Puromycin dihydrochloride | Sigma-Aldrich | Cat#P8833 |
| Q Sepharose™ Fast Flow | Cytiva | Cat#GE17-0510-10 |
| Recombinant Human SCF Protein | R&D Systems | Cat#255-SC |
| ReLeSR™ | Stemcell Technologies | Cat#100-0483 |
| RIPA Lysis and Extraction Buffer | Thermo Fisher Scientific | Cat#89900 |
| Sodium bicarbonate | Sigma-Aldrich | Cat#S5761 |
| Sodium bicarbonate | Sigma-Aldrich | Cat#1370131000 |
| SP Sepharose™ Fast Flow | Cytiva | Cat#GE17-0729-10 |
| StemPro Accutase | Thermo Fisher Scientific | Cat#A1110501 |
| TAK-242 | MedChemExpress | Cat#HY-11109 |
| TFEB activator 1 | MedChemExpress | Cat#HY-135825 |
| Thioflavin T | Sigma-Aldrich | Cat#596200 |
| Tris(2-carboxyethyl) phosphine (TCEP) | Sigma-Aldrich | Cat#75259-1G |
| Tris-buffered saline (TBS) | Thermo Fisher Scientific | Cat#J62938.K3 |
| Triton X-100 | Sigma-Aldrich | Cat#X100 |
| Triton X-114 | Sigma-Aldrich | Cat#X114 |
| Trypsin 0.25% | Thermo Fisher Scientific | Cat#15050065 |
| Trypsin Mass Spec Grade | Promega | Cat#V5280 |
| Trypsin-EDTA (0.5%), no phenol red | Thermo Fisher Scientific | Cat#15400054 |
| Tween-20 | Carl Roth | Cat#9127.1 |
| Uranyl acetate | SERVA Electrophoresis GmbH | Cat# 77870.02 |
| Value Fetal Bovine Serum | Thermo Fisher Scientific | Cat#A5256701 |
| Vitronectin (VTN-N) Recombinant Human Protein | Thermo Fisher Scientific | Cat#A400457 |
| Wheat germ agglutinin (WGA) | Thermo Fisher Scientific | Cat#W11261 |
| X-VIVO® 15 Serum-free Hematopoietic Cell Medium | Lonza Biosciences | Cat#02-060Q |
| Y-27632 (Dihydrochloride) | Stemcell Technologies | Cat#72302 |
| **Critical commercial assays** |  |  |
| Glial-Mag magnetofection kit | OZ Biosciences | Cat#KGL0250 |
| Pierce LAL Chromogenic Endotoxin Quantification Kit | Thermo Fisher Scientific | Cat#88282 |
| Pierce™ BCA Protein Assay Kits | Thermo Fisher Scientific | Cat#23225 |
| PROTEOSTAT® Protein Aggregation Assay | ENZO | Cat# ENZ-51023 |
| SYTOX® Blue Dead Cell stain solution | Thermo Fisher Scientific | Cat#S11348 |
| **Deposited data** |  |  |
| Mass spectrometry data of primary mouse microglia | https://www.ebi.ac.uk/pride/ | PXD085675 |
| **Experimental models: Cell lines** |  |  |
| KOLF2.1J iPSC | Jackson Laboratories | CVCL_B5P3; Cat#JIPSC001000 |
| **Experimental models: Organisms/strains** |  |  |
| Mouse: C57BL/6N | Charles River Laboratories | IMSR: JAX 000664 |
| **Oligonucleotides** |  |  |
| Negative control siRNA | Thermo Fisher Scientific | Cat#AM4641 |
| **SQSTM1 mouse siRNA** | Thermo Fisher Scientific | Cat#AM16708 |
| **Recombinant DNA** |  |  |
| human Alpha-synuclein | Addgene | Cat# #51437; RRID: Addgene_51437 |
| human Tau P301S (1N4R) | gift from Dr. Christina Ising DOI: 10.1038/s41586-019-1769-z | N/A |
| **Software and algorithms** |  |  |
| aPEAR, v 1.0.0 | VUGENE, LLC | N/A |
| FACSDIVA™ software, v9.0 | BD Biosciences | RRID: SCR_001456 |
| Fiji ImageJ, v2.0.0-rc-69/1.52n | Wayne Rasband | RRID: SCR_002285 |
| FlowJo, v3.05470 | FlowJo, LLC | RRID: SCR_008520 |
| Graph Pad Prism, v8.0 and v9.0 | GraphPad Software Inc. | RRID: SCR_002798 |
| Image Studio, v5.2 | LI-COR Biosciences | RRID: SCR_015795 |
| Imaris, v9.2.1 | Bitplane by Oxford Instruments plc | RRID: SCR_007370 |
| limma, v 3.60.6 | Bioconductor | RRID: SCR_010943 |
| R, v 4.4.0 | R Foundation for Statistical Computing | RRID: SCR_001905 |
| WebGestaltR, v 0.4.6 | WebGestaltR development team (Zhang Lab) | RRID: SCR_024312 |
| **Other** |  |  |
| 100 mesh formvar coated copper grid | Science Services | Cat#EFCF100-Cu-25 |
| 6-well plate | Falcon | Cat#353046 |
| Amicon® Ultra Zentrifugenfilter, 10 kDa MWCO | Merck Millipore | Cat#UFC9010 |
| BD FACSCANTOII | BD Biosciences | RRID: SCR_018055 |
| Bioruptor | Diagenode | RRID: SCR_023470 |
| C18-SD StageTips | CDS Empore | Cat#70-2019-1042-6EA |
| JEM-2100 Plus Transmission Electron Microscope | JEOL | RRID: SCR_020173 |
| McIlwain Tissue Chopper | Campden Instruments | Cat#TC752 |
| Millicell Cell Culture Insert, 30 mm, hydrophilic PTFE, 0.4 µm | Merck Millipore | Cat#PICM0RG50 |
| NanoDrop 2000c Spectrophotometer | Thermo Fisher Scientific | RRID: SCR_020309 |
| NuPAGE™ Bis-Tris Midi-Protein-Gele | Thermo Fisher Scientific | Cat#WG1402BOX |
| NuPAGE™ Bis-Tris Mini Protein Gels | Thermo Fisher Scientific | Cat#NP0335BOX |
| ODYSSEY CLx Imaging System | LI-COR Biosciences | RRID: SCR_014579 |
| Olympus SpinSR Microscope | Olympus | N/A |
| ReproSil-Pur column | Evosep | Cat#EV1109 Performance Column |
| SPARK Multimode Microplate reader | Tecan | RRID: SCR_021897 |
| T75 culture flasks | Greiner Bio-one | Cat#C7231 |
| TCS SP8 X Laser Scanning Microscope | Leica | N/A |
| Trans-Blot Turbo Midi 0.2 µm Nitrocellulose Transfer Packs | Bio-Rad | Cat#1704159 |
| Trans-Blot Turbo Mini 0.2 µm Nitrocellulose Transfer Packs | Bio-Rad | Cat#1704158 |
| Vibratome VT1200S | Leica | RRID: SCR_018453 |

### Experimental models and subject details

#### Ethical considerations

Animal procedures were conducted in accordance with the institutional guidelines of the Max Planck Society and the European Directive 2010/63/EU. The use of animals for tissue collection was registered with and acknowledged by the competent local veterinary authority (Veterinäramt; registration no. MPIa_Anzeige§4_SB.24.01 and MPIa_Anzeige§4_SB.26.02). Human postmortem brain tissue was obtained from the DZNE Brain Bank (German Center for Neurodegenerative Diseases, Bonn, Germany). The use of human tissue for this study was approved by the Ethics Committee of the University of Cologne (approval no. 24-1176). Written informed consent for brain donation and the use of tissue and associated clinical information for research purposes was obtained from the donors or their legal representatives in accordance with DZNE Brain Bank procedures. All procedures involving human tissue were conducted in accordance with the Declaration of Helsinki.

#### Animal procedures

All mice were on a C57BL/6N genetic background. Mice were housed under standard conditions at 22 °C and a 12 hrs light-dark cycle with free access to food and water. Animal care and handling were performed according to the animal welfare guidelines laid down by the German Research Council (DFG) and approved by the local ethical committees.

#### Primary microglia cultures

Primary microglia cultures were isolated from neonatal wild-type C57BL/6N mice (P0-P3) as previously described^59^. In brief, pups were euthanized by decapitation, and the brains were rapidly removed. After careful removal of the meninges, brain tissue was dissociated using mechanical shearing and trypsin. Cells were seeded into poly-L-lysine (PLL, Sigma-Aldrich) coated T75 culture flasks (Greiner Bio-One) and maintained in Dulbecco’s Modified Eagle’s Medium (DMEM, Thermo Fisher Scientific) supplemented with 10% heat-inactivated fetal bovine serum (FBS; Thermo Fisher Scientific) and 1% penicillin/streptomycin (P/S; Thermo Fisher Scientific). After 7-10 days, mature microglia were shaken off the astrocytic layer. This harvesting procedure was repeated every other day for up to three rounds. Isolated microglia were seeded according to the experiment and allowed to adhere overnight in DMEM 1%P/S and 1X *N*-2 supplement (Thermo Fisher Scientific).

#### iPSC culture

KOLF2.1J-CSF1R-HaloTag iPSCs (Jackson Laboratories) were maintained on vitronectin-coated 6-well plates in mTeSR+ medium (Stemcell Technologies) at 37 °C, 5% CO₂, and 100% humidity. At approximately 80% confluency, cells were passaged using ReLeSR (Stemcell Technologies). Briefly, cells were washed with Ca²⁺/Mg²⁺-free PBS, incubated with ReLeSR for 5-6 min at 37 °C, and collected as small clumps using wash buffer (Advanced DMEM/F12 supplemented with 0.1% BSA; Thermo Fisher Scientific). Following centrifugation at 300 × g for 5 min at room temperature, cells were replated on vitronectin-coated plates in mTeSR+ supplemented with 10 µM Y-27632 (Stemcell Technologies) for the first 24 h. Medium was subsequently changed every other day.

#### Embryoid body-based iMic differentiation

iPSCs were differentiated into microglial progenitors using a modified embryoid body (EB)-based protocol^60^. iPSCs were dissociated into single cells using Accutase (Thermo Fisher Scientific) for 5-6 min, collected in wash buffer, centrifuged at 300 × g for 5 min at room temperature, and counted. For EB formation, 3 × 10⁶ cells were seeded per well of an anti-adherence-treated AggreWell800 24-well plate (Stemcell Technologies) in 2 ml EB induction medium (mTeSR+ supplemented with 20 ng/ml SCF, 50 ng/ml BMP4, and 50 ng/ml VEGF; Miltenyi Biotec). Y-27632 (10 µM) was added for the first 24 h. Plates were centrifuged at 800 × g for 3 min without braking, and EBs were cultured at 37 °C, 5% CO₂, and 100% humidity with daily 75% medium changes.

After 5 days, EBs were transferred to two 6-well plates containing 2 ml EB differentiation medium per well (X-VIVO 15, 2 mM GlutaMAX, 0.55 mM β-mercaptoethanol, 100 U/ml penicillin, 100 µg/ml streptomycin, 25 ng/ml IL-3, and 100 ng/ml CSF1). Cultures were maintained under the same conditions with complete medium changes every 7 days. From approximately day 21, non-adherent microglial progenitors were collected during medium changes, filtered through a 70 µm cell strainer to retain EBs, and centrifuged at 300 × g for 5 min. After counting, progenitors were resuspended in EB differentiation medium at 10,000 cells/µl for engraftment into organotypic slice cultures.

#### Generation of chimeric organotypic slice cultures

Chimeric organotypic slice cultures (cOSCs) were generated as described elsewhere^61^, with minor modifications. Hippocampi from P4-P6 C57BL/6N mice were isolated in cold preparation medium (phenol red-free MEM; Thermo Fisher Scientific) and sectioned into 350 µm slices using a McIlwain tissue chopper. Intact slices were transferred onto Millicell culture inserts (Merck Millipore; 3-4 slices/insert) in 6-well plates containing pre-warmed slice culture medium (MEM supplemented with 20% horse serum, 1 mM GlutaMAX, 0.00125% ascorbic acid, 0.001 mg/ml insulin, 1 mM CaCl₂, 2 mM MgSO₄, 13 mM D-glucose, and 100 U/ml penicillin/100 µg/ml streptomycin; pH 7.28) and 5 µg/ml anti-mouse CSF1R antibody (CD115; BioLegend) for endogenous microglia depletion.

After 4 days, human iPSC-derived microglial progenitors were harvested as described above, and 10,000 cells in 1 µl EB differentiation medium were seeded onto each slice. Cultures were maintained for an additional 3 weeks to allow microglial differentiation and engraftment before experiments. Cultures were kept at 37 °C, 5% CO₂, and 100% humidity, with medium changes three times per week and replenishment of the depletion antibody at each change.

### Methods Details

#### Protein Purification

Human WT α-Syn and Tau P301S (1N4R) was expressed in Escherichia coli BL21 (DE3) CodonPlus cells transformed with the human α-Syn (gift from Michael J Fox Foundation MJFF, Addgene plasmid #51437) or Tau P301S (1N4R) (gift from Dr. Christina Ising) expression plasmid. Protein expression was induced with 0.6 mM IPTG at an OD600 of 0.8 for 3.5 hrs at 37 °C. Bacterial pellets were lysed by sonication in BRB-80 buffer containing protease inhibitors and 0.1% β-mercaptoethanol. Following the removal of the precipitate, the lysate was heat-treated (30 min) to precipitate bacterial proteins, and the soluble fraction was purified by cation-exchange chromatography using Q or SP Sepharose Fast Flow resin (Cytiva) with stepwise NaCl elution (0-1 M). Fractions containing α-Syn or Tau were identified by SDS-PAGE/Coomassie staining, pooled, buffer-exchanged and concentrated using 10-kDa centrifugal filters (Merck Millipore) into PBS. Protein concentration was determined by BCA assay (Thermo Fisher Scientific), and purified protein was aliquoted and stored at −80°C until further use. All preparations were cleaned as described elsewhere by Triton-X114 phase separation^62^. Afterwards endotoxin levels were quantified using the Pierce LAL Chromogenic Endotoxin Quantification Kit (Thermo Fisher Scientific) according to manufacturer’s protocol. Endotoxin levels below 0.01 EU/µg were considered as endotoxin free. Finally, purified proteins were assembled into PFFs as described previously and after labelled with ATTO-405 NHS-ester, ATTO-488 NHS-ester, or ATTO-647N NHS-ester (Atto-Tec GmbH) fluorophore according to manufacturer’s protocol^63^. The fibrillar nature of PPFs were monitored by Thioflavion T assay and visualized by Transmission Electron Microscopy (TEM). PFFs were fragmented by sonication as described previously to fragments of 30-120 nm^63^.

#### Electron microscopy

α-Syn and Tau fibrils and their subsequent PFFs were fixed with 1% formaldehyde (Science Services). A 100 mesh formvar coated copper grid (Science Services) was placed onto 5 µl of sample and incubated for 20 min. Grids were washed seven times for 2 min in PBS and placed for 5 min on a drop of 1% glutaraldehyde (Merck Millipore). Grids were washed eight times in ddH_2_O. Grids were placed for four min onto a drop of 1.5% aqueous uranyl acetate and blotted with a filter paper. Images were acquired using a JEM-2100 Plus Transmission Electron Microscope (JEOL) operating at 80 kV equipped with a OneView 4K camera (Gatan).

#### NF-κB nuclear translocation assay

Microglia were seeded at a density of 4 × 10⁴ cells/well in 8-well µ-Slides (ibidi). Cells were treated with or without LPS (10 ng/ml) in the presence or absence of TAK-242 (500 nM), CHX (50 µg/mL), or TAc1 (2 µM). To assess the initial induction of NF-κB nuclear translocation, cells were fixed after 30 min of treatment. To investigate the persistence of NF-κB nuclear localization, cells were fixed 24 h after treatment washout. Cells were subsequently immunostained for NF-κB as described in the immunocytochemistry section.

#### Puromycin incorporation assay

Microglia were seeded at a density of 1 × 10⁶ cells/well. For TLR4 inhibition experiments, cells were treated with or without LPS (10 ng/ml) in the presence or absence of TAK-242 (500 nM) for 24 h. For translation inhibition experiments, cells were treated with or without LPS (50 ng/ml) and CHX (50 µg/mL) for 2 h, followed by washout and a 24 h recovery period. To assess nascent protein synthesis, puromycin (1 µM) was added during the final hour of culture. Cells were subsequently harvested, and cell pellets were collected for Western blot analysis of puromycin incorporation.

#### Uptake and Degradation experiments

For all uptake and degradation experiments microglia were plated at a density of 2.5 x 10^5^ cells/cm^2^ and treated with 1 µM α-Syn PFFs and 10 ng/ml LPS (InvivoGen) in a total volume of 1 ml/1 mio cells unless otherwise stated. Afterwards, α-Syn PFFs were washed away once with PBS, replaced by fresh DMEM 1% P/S and 1X *N*-2 supplement and followed up for 24 - 72 hrs. Furthermore, cells were treated with CHX (50 µM), TAK-242 (500 nM), TAc1 (2 µM), Puromycin (1 µM) and Bafilomycin A1 (100 nM) as indicated in the corresponding figure schematics. Treatment concentrations and durations are provided in the respective scheme or figure legends.

#### PROTEOSTAT® staining

Intracellular protein aggregation was assessed using the PROTEOSTAT® Protein Aggregation Assay (ENZO). Microglia were seeded at a density of 4 × 10⁴ cells/well in 8-well µ-Slides (ibidi) and were either left untreated or treated with AZC (1 mM) for 24 h. Following treatment, cells were fixed with 4% PFA, permeabilized with PBST, and stained with PROTEOSTAT® dye (1:10,000 for 30 min).

#### Magnetofection

Primary microglia were magnetofected 24 hrs after seeding using the Glial-Mag magnetofection kit (OZ Biosciences) according to the manufacturer’s protocol. In brief, siRNA (non-targeting control pool (scramble or SQSTM1)) was diluted in OPTIMEM (Thermo Fischer Scientific) and mixed with Glial-Mag transfection reagent. After 20 min incubation at room temperature, the solution was added to each respective well in a dropwise manner. The Glia-Boost reagent was added directly to cells to a final concentration of 1X. The final concentration of siRNA was 25 nM. Culture plates were kept on top of the magnetic plate for 30 min and medium was replaced after 3 hrs. Kock-down efficiency was analysed after 48 hrs by western blot.

#### Flow cytometry

For uptake and degradation experiments cells were lifted with Trypsin 0.5%/EDTA 10X (Thermo Fisher Scientific) and collected in tubes containing FACS buffer (PBS, 2% FCS, 1 µg/ml DNase I, 1X SYTOX dead cell stain). Samples were stored on ice and analysed immediately on a BD FACS Canto II using BD DIVA Software. Further analysis was carried out in FlowJO.

#### Proteomics

##### Treatment of cells

Primary microglia were seeded in 6-well plates at a density of 1 × 10^6^ cells per well and allowed to adhere overnight. Cells were then treated with LPS (10 ng/ml), α-Syn PFFs (1 µM), or a combination of LPS and α-Syn PFFs. After 24 h, cells were washed with PBS, detached by trypsinization, and collected by centrifugation at 300 × g for 5 min.

##### Protein extraction, digestion, and peptide desalting

Cell pellets were lysed in 6 M guanidinium hydrochloride, 100 mM Tris-HCl containing 2.5 mM tris(2-carboxyethyl)phosphine (TCEP) and 10 mM chloroacetamide (CAA), followed by heating (95°C, 10 min), sonication, and centrifugation (20k x g, 20 min). Protein concentrations were determined by NanoDrop after dilution in 20 mM Tris-HCl. Up to 100 μg of protein was diluted 10-fold and digested overnight at 37°C with sequencing-grade trypsin (Promega) at an enzyme-to-protein ratio of 1:200 (w/w). On the next day, digestion was terminated by addition of formic acid to a final concentration of 1%. Peptides were desalted using C18 StageTips (Empore, 3M), dried by vacuum centrifugation, and reconstituted in 0.1% formic acid prior to LC-MS/MS analysis.

The Evosep One liquid chromatography system^64^ was used for analysing the samples with the predefined 60 samples per day (60SPD) method. The analytical column we used was an ReproSil-Pur column, 8 cm x 150 µm, with 1.5 µm C18 beads (EV1109 Performance Column, Evosep). The mobile phases A and B were 0.1 % formic acid in water and 0.1% formic acid in 100% ACN, respectively.

##### Mass spectrometry

Peptides were analyzed on a hybrid TIMS quadrupole TOF mass spectrometer (timsTOF HT, Bruker) in a data-independent acquisition parallel accumulation, serial fragmentation (diaPASEF) mode. The mass spectra range was set to 350 - 1200 m/z and TIMS ion accumulation and ramp times were set to either 50 ms and total cycle time was 0.73 s. The ion mobility range was set to 1/K0 = 0.8 - 1.25 V-s/cm2. Isolation windows in the m/z versus ion mobility plane were defined to cover the region of highest precursor ion density using py DIA^65^. Collision energy was applied linearly with ion mobility from 0.6 to 1.6 V-s/cm2, and collision energy from 20 to 59 eV.

##### Data Analysis

Raw data was analyzed using Spectronaut (v20.0.25, Biognosys) using the default parameters against the one-protein-per-gene reference proteome for *Mus musculus*, UP000000589, downloaded August, 2022. Methionine oxidation and protein N-terminal acetylation were set as variable modifications; cysteine carbamidomethylation was set as fixed modification. The digestion parameters were set to “specific” and “Trypsin/P”, with two missed cleavages permitted. Protein groups were filtered for at least two valid values in at least one comparison group and missing values were imputed from a normal distribution with a down-shift of 1.8 and standard deviation of 0.3. Differential expression analysis was performed using limma (v3.60.6) in R (v 4.4.0)^66^. Gene set enrichment analysis (GSEA) was performed using WebGestaltR (v0.4.6)^67^. Clustering and visualization of the GSEA results was done using aPEAR (v1.0.0)^68^.

### Immunocytochemistry

Cultures were fixed in 4% PFA dissolved in DPBS (Thermo Fisher Scientific) and permeabilized using PBS containing 0.1% Triton X-100 (PTX). A blocking solution containing PTX and 5% normal goat serum (NGS, Abcam) or normal donkey serum (NDS, Jackson Immuno Research) was applied for 30 min. After this, sections were incubated in mixtures of primary antibodies against NFκB p65 (rabbit, 1:1,000, Cell Signaling Technology), p62/SQSTM1 (rabbit, 1:2,000, Cell Signaling Technology), and Cathepsin D (goat, 1:1,000, Thermo Fisher Scientific) overnight at 4°C. After incubation, sections were washed in PTX and incubated overnight at 4°C in the mixture of secondary antibodies, all diluted in blocking solution. DAPI was used for nuclear counterstain and Phalloidin coupled to AlexaFluor-488 or TexasRed at 0.05 mg/mL and 0.33 µM for 20 min in PBS. Confocal images were acquired using a Leica TCS SP8 confocal microscope equipped with the LIGHTNING super-resolution module (Leica Microsystems). For quantitative comparisons, all samples within an experiment were imaged using identical acquisition settings, including laser power, detector gain, and offset. Images were processed using Leica LAS X software with the LIGHTNING reconstruction algorithm, applying identical processing parameters to all samples within each experiment.

Images were processed in Fiji (ImageJ) or Imaris (v9.2.1). A custom Fiji macro was used to automatically identify and quantify α-Syn PFFs, lysosomes or p62-condensates based on thresholding and particle analysis. The same processing parameters were applied to all images within each experiment.

### Western Blot

Samples were lysed in 1X RIPA buffer (Thermo Fisher Scientific) containing 1X HALT Protease inhibitor cocktail (Thermo Fisher Scientific) and phophoSTOP (Sigma-Aldrich) for 20 min on ice. The soluble fraction was collected by centrifugation at 16.000 xg for 10 min. Samples were denatured in 1X Laemmli Buffer containing SDS and heated for 5 min to 95 °C. Samples were run on NuPAGE 4-12% Bis-Tris gels (Thermo Fisher Scientific) in MES buffer (Thermo Fisher Scientific) at 150 V for one hour. Afterwards proteins were transferred to NitroCellulose Membranes using a semi-dry TurboBlot System (Bio-Rad). Membranes were blocked in Tris-buffered saline containing 0.1% Tween20 (TBST) and 5% bovine serum albumin (BSA) for 30 min. Afterwards primary antibodies against puromycin (rabbit, 1:1,000, Cell Signaling Technology), α-Syn (mouse, 1:1,000, BioLegends), beta actin (mouse, 1:2,000, Proteintech), MAP1LC3b (rabbit, 1:1,000, Sigma-Aldrich), p62/SQSTM1 (rabbit, 1:2,000, Cell Signaling Technology), and Cathepsin D (goat, 1:1,000, Thermo Fisher Scientific were applied over night at 4 °C, followed by goat anti-rabbit, goat anti-mouse secondary IRdye antibodies (700 or 800, (LiCOR Bioscience)) or donkey anti-goat IgG (H + L) Alexa Fluor 647 (Thermo Fisher Scientific) for 1 hour at room temperature. Membranes were analysed using a LiCOR system operated with ImageStudio (v5.0).

### Treatment of cOSC with α-Syn and LPS

cOSCs were treated with 2 µM Atto 647-labelled α-Syn and/or 200 ng/ml LPS for 48 h. Subsequently, the medium was replaced with regular slice culture medium containing the depletion antibody for an additional 48 h to allow α-Syn degradation. Slices were then washed with PBS, fixed with 4% PFA in PBS for 2 h at room temperature, washed three times with PBS, and stored in PBS at 4 °C until further processing.

### Immunostaining of cOSC

Individual cOSCs were carefully transferred from culture inserts onto microscopy slides using a fine brush, preserving tissue orientation, and air-dried for 1 h at room temperature. Slices were rehydrated in PBS for 15 min and blocked with 5% NDS and 0.3% Triton X-100 in PBS for 2 h at room temperature. Primary antibodies against Cathepsin D (goat, 1:500; Thermo Fisher Scientific) and Iba1 (rabbit, 1:250; Wako) were incubated overnight at 4 °C in PBS containing 2% NDS and 0.3% Triton X-100. Following three PBS washes, slices were incubated with DAPI and Alexa Fluor-conjugated donkey secondary antibodies (Thermo Fisher Scientific) in PBS containing 1% NDS and 0.3% Triton X-100 for 2 h at room temperature. After three additional PBS washes, slices were mounted with Immu-Mount (Epredia), dried overnight at room temperature, and imaged using an Olympus SpinSR microscope equipped with a 60x silicone oil-immersion objective.

### Immunofluorescence staining of human brain tissue

Human brain tissue (frontal cortex) was obtained as fresh-frozen tissue blocks from the DZNE Brain Bank. Tissue was thawed overnight in 4% PFA at 4 °C, followed by an additional 2 hrs fixation at room temperature. After extensive washing in PBS (3 x 30 min), tissue was sectioned into 50-µm-thick free-floating sections using a vibratome. Sections were permeabilized in PBS containing 0.3% Triton X-100 (PBST) supplemented with 0.3% saponin for 1 hr, followed by blocking for 1 hr in PBST/0.3% saponin containing 3% BSA and 3% NDS. Sections were then incubated with primary antibodies against α-Syn pSer129 (mouse, 1:500, BioLegend), Cathepsin D (goat, 1:500; Thermo Fisher Scientific) and Iba1 (rabbit, 1:50; Wako) overnight at 4 °C. Following washing in PBST, sections were incubated with the appropriate fluorophore-conjugated secondary antibodies for 2 hrs at RT in PBST/0.3% saponin containing 3% BSA and 3% NDS. Nuclei were counterstained with DAPI before sections were mounted onto glass slides using ProLong™ Diamond Antifade Mountant (Thermo Fisher Scientific). Images were acquired using a Leica SP8-X laser-scanning confocal microscope equipped with a 60x oil-immersion objective.

### Statistical analysis

Each n represents an independent biological experiment. Data were analyzed using GraphPad Prism (v.10) and presented as mean ± SEM of at least three independent experiments. Data were tested for Gaussian distribution (Shapiro-Wilk or D’Agostino-Pearson test). Where data passed the normality test, control versus treatment comparisons were analyzed with an unpaired two-tailed Student’s t-test (two groups) or a one-way ANOVA with Tukey’s post hoc test (≥3 groups). Non-normally distributed data were analyzed with the Kruskal-Wallis test with Dunn’s post hoc test. Proteomic datasets were analyzed using limma with false discovery rate correction as described in the Proteomics section. Levels of significance are indicated as *p < 0.05; **p < 0.01; ***p < 0.001; ****p < 0.0001; ns, not significant.

