## Supplementary material for "Inflammatory proteostasis reprogramming creates a lysosomal bottleneck for microglial aggregate clearance": Figure S1

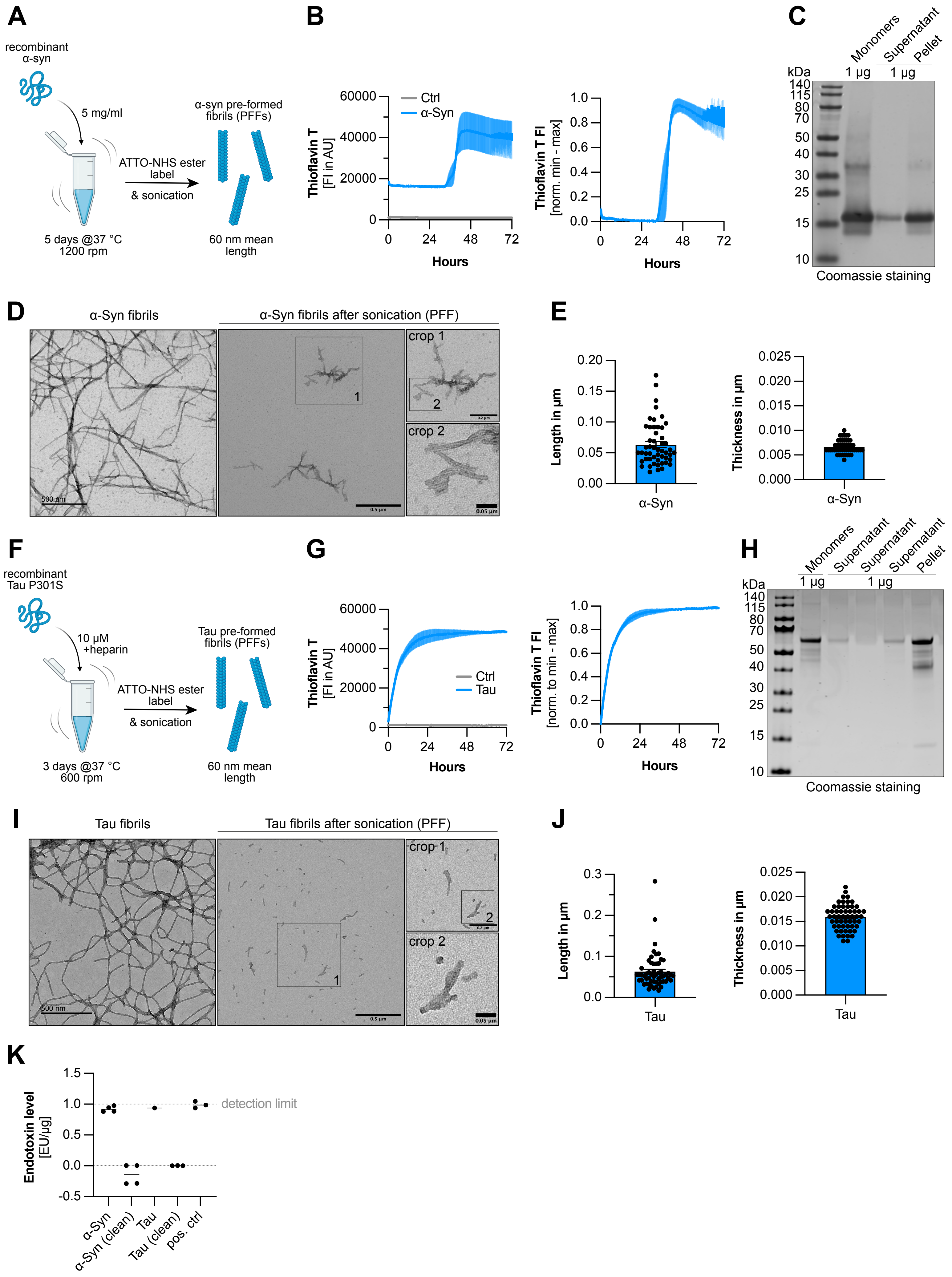
A
recombinant α-syn
5 mg/ml
ATTO-NHS ester label
& sonication
α-syn pre-formed fibrils (PFFs)
5 days @37 °C 1200 rpm
60 nm mean length
B
Ctrl
α-Syn
Thioflavin T [FI in AU]
Thioflavin T FI [norm. min - max]
Hours
C
Monomers
Supernatant
Pellet
1 µg
1 µg
kDa
Coomassie staining
D
α-Syn fibrils
α-Syn fibrils after sonication (PFF)
crop 1
crop 2
500 nm
0.5 µm
0.2 µm
0.05 µm
E
Length in µm
Thickness in µm
α-Syn
F
recombinant Tau P301S
10 µM +heparin
ATTO-NHS ester label
& sonication
Tau pre-formed fibrils (PFFs)
3 days @37 °C 600 rpm
60 nm mean length
G
Ctrl
Tau
Thioflavin T [FI in AU]
Thioflavin T FI [norm. to min - max]
Hours
H
Monomers
Supernatant
Supernatant
Supernatant
Pellet
1 µg
1 µg
kDa
Coomassie staining
I
Tau fibrils
Tau fibrils after sonication (PFF)
crop 1
crop 2
500 nm
0.5 µm
0.2 µm
0.05 µm
J
Length in µm
Thickness in µm
Tau
K
Endotoxin level [EU/µg]
detection limit
α-Syn
α-Syn (clean)
Tau
Tau (clean)
pos. ctrl
