## Supplementary material for "Inflammatory proteostasis reprogramming creates a lysosomal bottleneck for microglial aggregate clearance": Figure S2

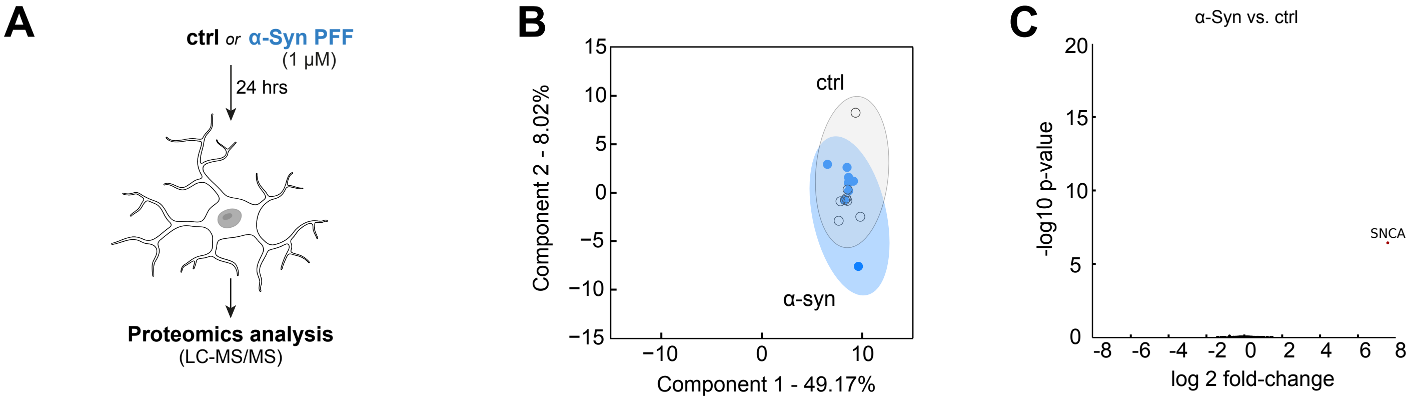
A
ctrl or α-Syn PFF
(1 μM)
24 hrs
Proteomics analysis
(LC-MS/MS)
B
ctrl
α-syn
Component 2 - 8.02%
Component 1 - 49.17%
C
α-Syn vs. ctrl
SNCA
-log10 p-value
log 2 fold-change
