## Supplementary material for "Inflammatory proteostasis reprogramming creates a lysosomal bottleneck for microglial aggregate clearance": Figure S3

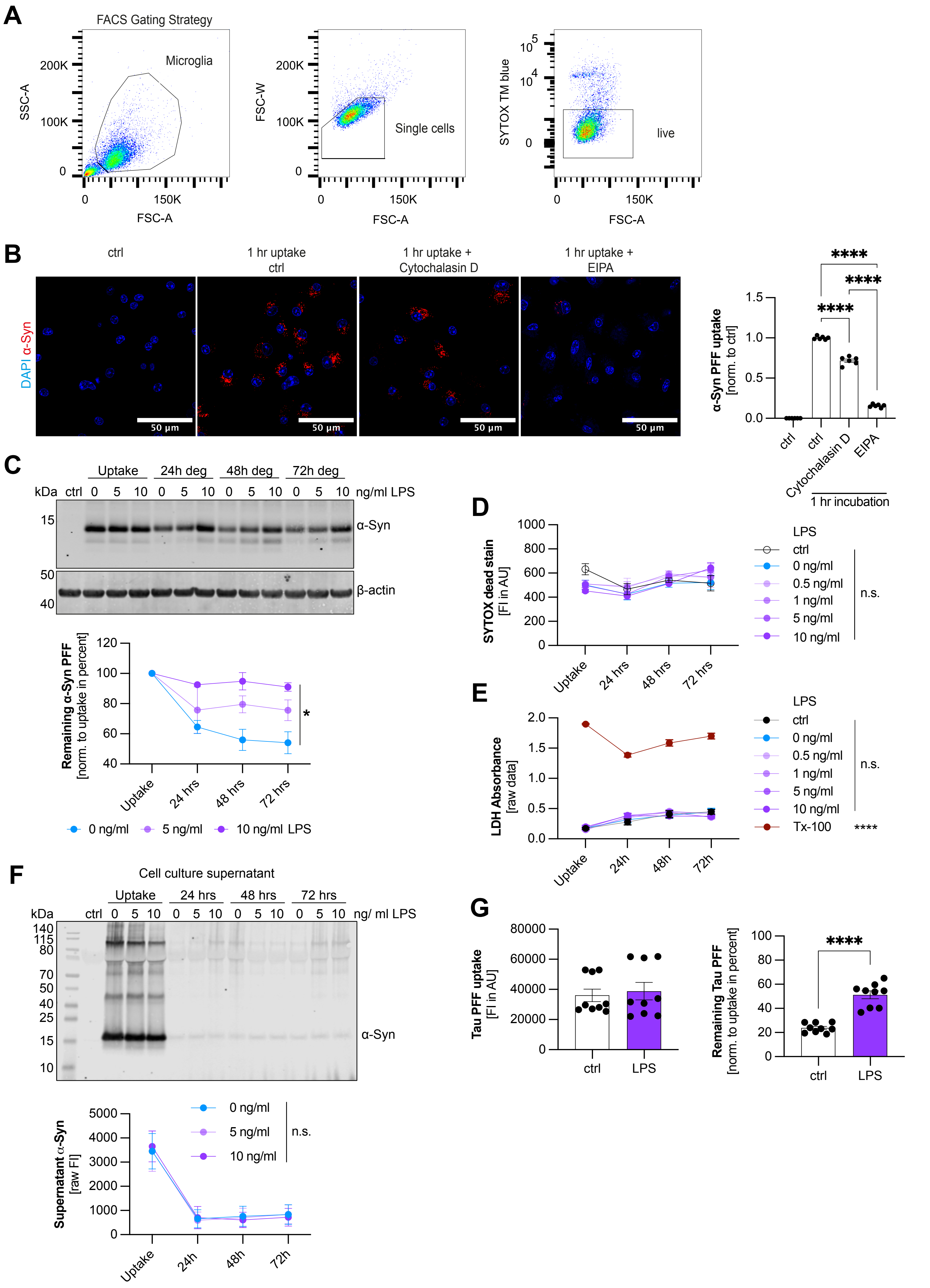

A
FACS Gating Strategy
Microglia
SSC-A
FSC-A
FSC-W
Single cells
SYTOX TM blue
live
B
ctrl
1 hr uptake ctrl
1 hr uptake + Cytochalasin D
1 hr uptake + EIPA
DAPI α-Syn
50 µm
α-Syn PFF uptake [norm. to ctrl]
ctrl
Cytochalasin D
EIPA
1 hr incubation
****
C
Uptake
24h deg
48h deg
72h deg
kDa
ng/ml LPS
α-Syn
β-actin
Remaining α-Syn PFF [norm. to uptake in percent]
24 hrs
48 hrs
72 hrs
0 ng/ml
5 ng/ml
10 ng/ml LPS
*
D
SYTOX dead stain [FI in AU]
LPS
0.5 ng/ml
1 ng/ml
10 ng/ml
n.s.
E
LDH Absorbance [raw data]
Tx-100
F
Cell culture supernatant
24 hrs
48 hrs
72 hrs
ng/ ml LPS
Supernatant α-Syn [raw FI]
24h
48h
72h
G
Tau PFF uptake [FI in AU]
LPS
Remaining Tau PFF [norm. to uptake in percent]
