## Supplementary material for "Inflammatory proteostasis reprogramming creates a lysosomal bottleneck for microglial aggregate clearance": Figure S4

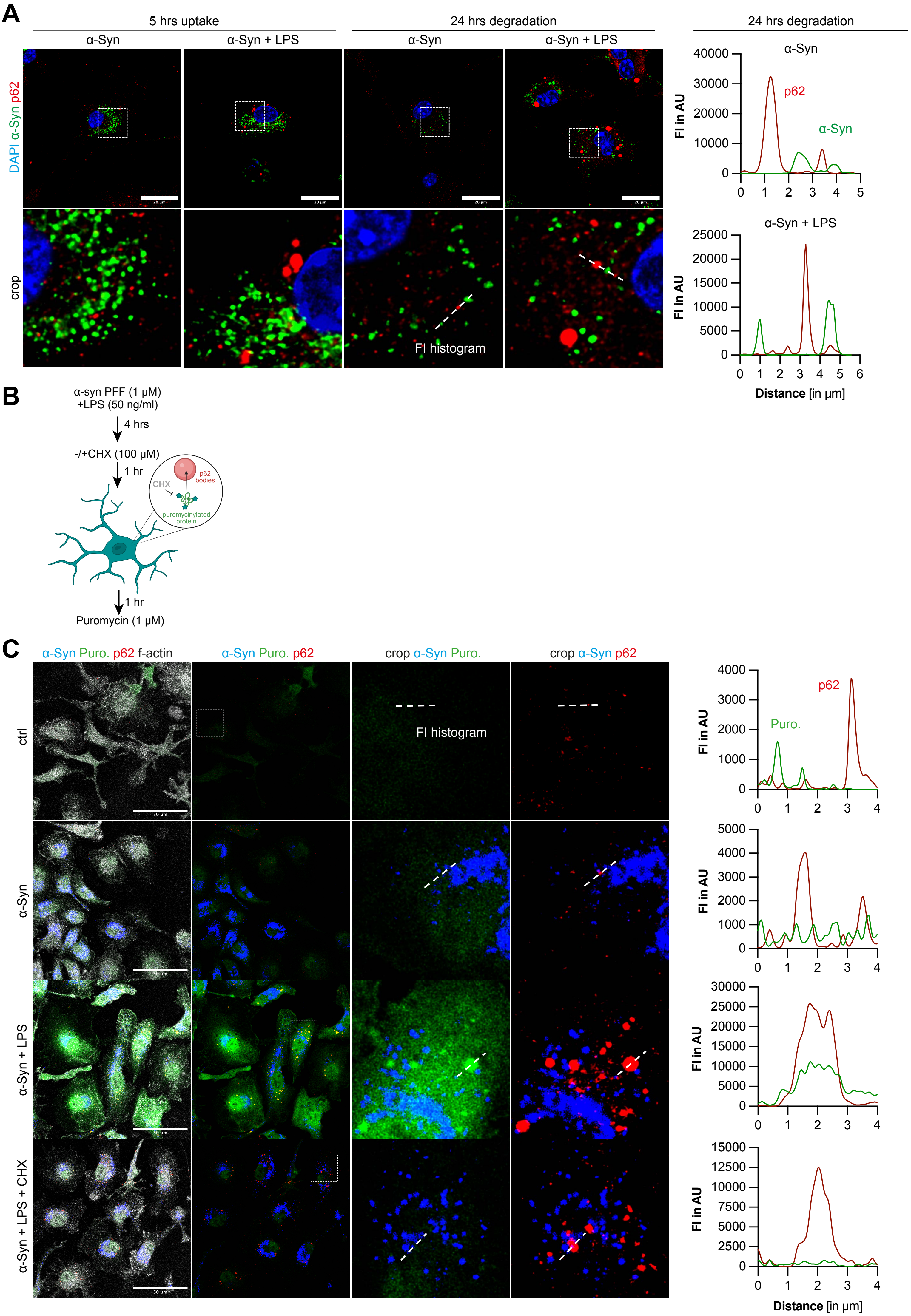
A
5 hrs uptake
24 hrs degradation
α-Syn
α-Syn + LPS
DAPI α-Syn p62
crop
20 µm
FI histogram
24 hrs degradation
α-Syn
p62
α-Syn
FI in AU
40000
30000
20000
10000
0
0 1 2 3 4 5
α-Syn + LPS
25000
20000
15000
10000
5000
0
0 1 2 3 4 5 6
Distance [in µm]
B
α-syn PFF (1 µM)
+LPS (50 ng/ml)
4 hrs
-/+CHX (100 µM)
1 hr
CHX
p62 bodies
puromycinylated protein
1 hr
Puromycin (1 µM)
C
α-Syn Puro. p62 f-actin
α-Syn Puro. p62
crop α-Syn Puro.
crop α-Syn p62
ctrl
α-Syn
α-Syn + LPS
α-Syn + LPS + CHX
50 µm
FI histogram
p62
Puro.
FI in AU
4000
3000
2000
1000
0
0 1 2 3 4
5000
4000
3000
2000
1000
0
30000
25000
20000
15000
10000
5000
0
15000
12500
10000
7500
5000
2500
0
Distance [in µm]
