## Supplementary material for "Inflammatory proteostasis reprogramming creates a lysosomal bottleneck for microglial aggregate clearance": Figure S5

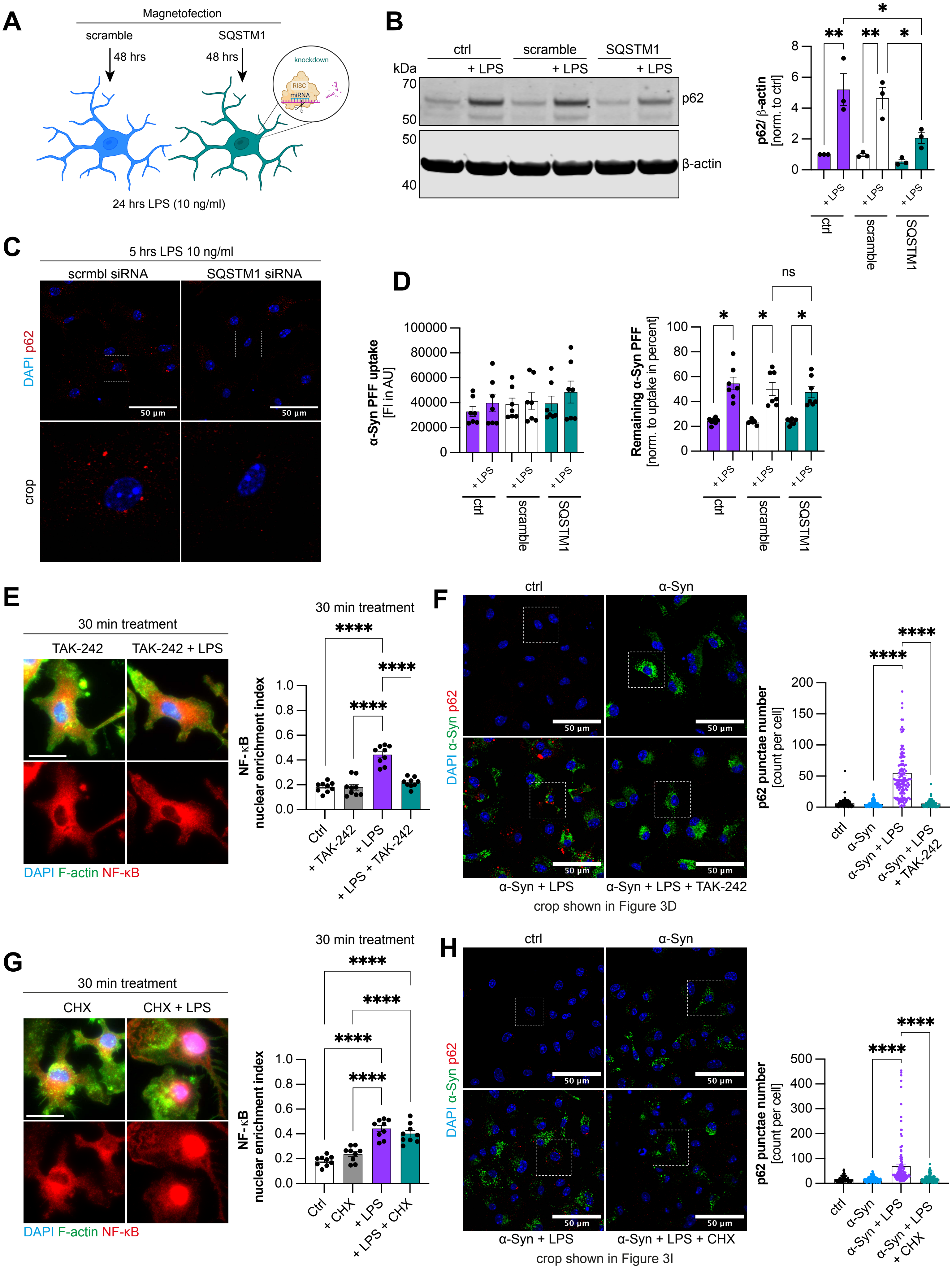

A
Magnetofection
scramble
SQSTM1
48 hrs
48 hrs
knockdown
RISC
miRNA
24 hrs LPS (10 ng/ml)
B
ctrl
scramble
SQSTM1
kDa
+ LPS
+ LPS
+ LPS
70
50
50
40
p62
β-actin
p62/ β-actin
[norm. to ctrl]
C
5 hrs LPS 10 ng/ml
scrmbl siRNA
SQSTM1 siRNA
DAPI p62
crop
50 µm
D
α-Syn PFF uptake
[FI in AU]
Remaining α-Syn PFF
[norm. to uptake in percent]
ns
E
30 min treatment
TAK-242
TAK-242 + LPS
DAPI F-actin NF-κB
NF-κB
nuclear enrichment index
Ctrl
+ TAK-242
+ LPS
+ LPS + TAK-242
F
ctrl
α-Syn
DAPI α-Syn p62
α-Syn + LPS
α-Syn + LPS + TAK-242
crop shown in Figure 3D
p62 punctae number
[count per cell]
α-Syn + LPS
α-Syn + LPS + TAK-242
G
30 min treatment
CHX
CHX + LPS
DAPI F-actin NF-κB
+ CHX
+ LPS + CHX
H
α-Syn + LPS + CHX
crop shown in Figure 3I
α-Syn + LPS + CHX
