## Supplementary material for "Inflammatory proteostasis reprogramming creates a lysosomal bottleneck for microglial aggregate clearance": Figure S6

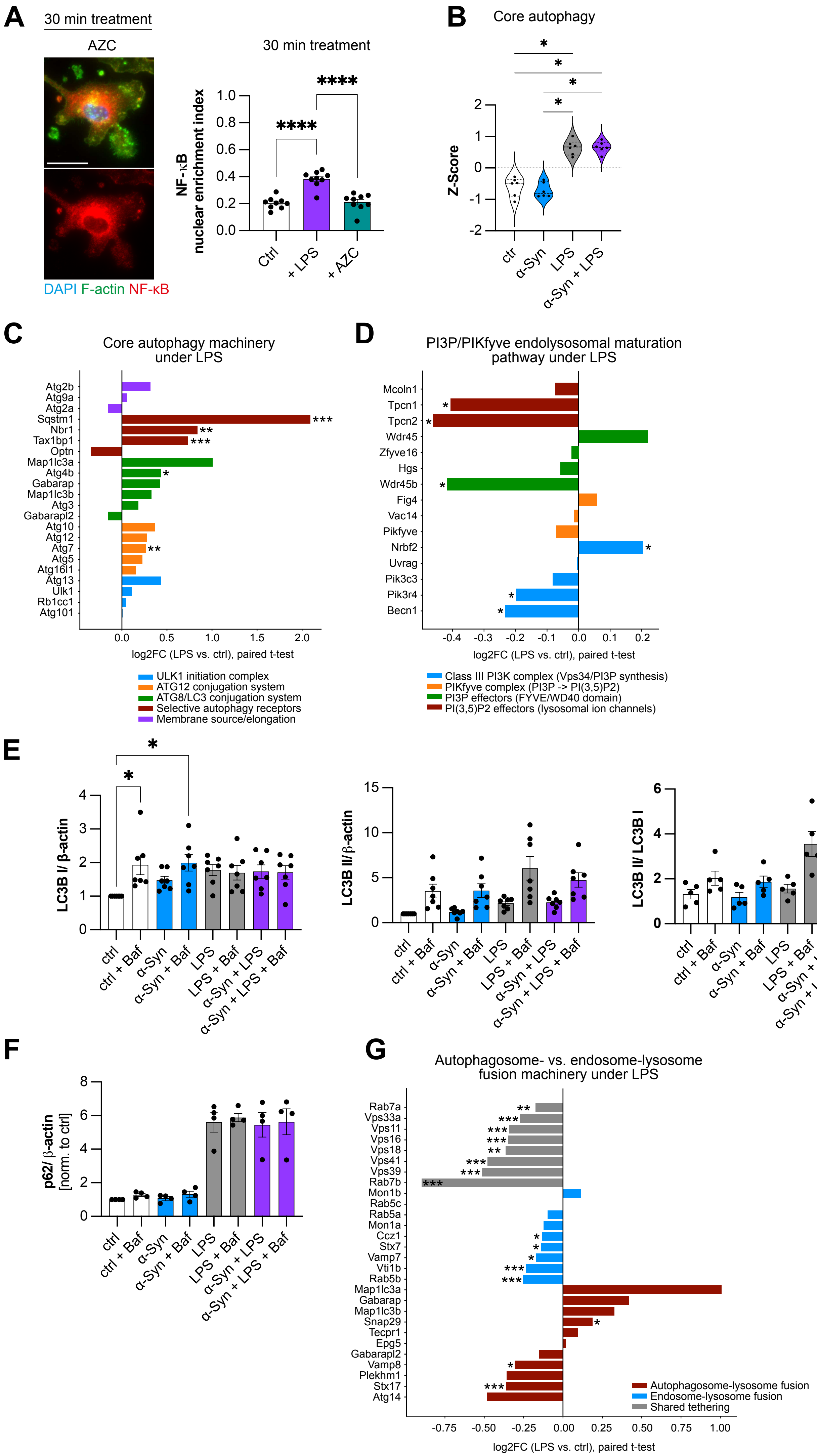

A
30 min treatment
AZC
DAPI F-actin NF-κB
30 min treatment
NF-κB nuclear enrichment index
Ctrl
+ LPS
+ AZC
B
Core autophagy
Z-Score
ctr
α-Syn
LPS
α-Syn + LPS
C
Core autophagy machinery under LPS
Atg2b
Atg9a
Atg2a
Sqstm1
Nbr1
Tax1bp1
Optn
Map1lc3a
Atg4b
Gabarap
Map1lc3b
Atg3
Gabarapl2
Atg10
Atg12
Atg7
Atg5
Atg16l1
Atg13
Ulk1
Rb1cc1
Atg101
log2FC (LPS vs. ctrl), paired t-test
ULK1 initiation complex
ATG12 conjugation system
ATG8/LC3 conjugation system
Selective autophagy receptors
Membrane source/elongation
D
PI3P/PIKfyve endolysosomal maturation pathway under LPS
Mcoln1
Tpcn1
Tpcn2
Wdr45
Zfyve16
Hgs
Wdr45b
Fig4
Vac14
Pikfyve
Nrbf2
Uvrag
Pik3c3
Pik3r4
Becn1
log2FC (LPS vs. ctrl), paired t-test
Class III PI3K complex (Vps34/PI3P synthesis)
PIKfyve complex (PI3P -> PI(3,5)P2)
PI3P effectors (FYVE/WD40 domain)
PI(3,5)P2 effectors (lysosomal ion channels)
E
LC3B I/ β-actin
LC3B II/β-actin
LC3B II/ LC3B I
ctrl
ctrl + Baf
α-Syn
α-Syn + Baf
LPS
LPS + Baf
α-Syn + LPS
α-Syn + LPS + Baf
F
p62/ β-actin [norm. to ctrl]
G
Autophagosome- vs. endosome-lysosome fusion machinery under LPS
Rab7a
Vps33a
Vps11
Vps16
Vps18
Vps41
Vps39
Rab7b
Mon1b
Rab5c
Rab5a
Mon1a
Ccz1
Stx7
Vamp7
Vti1b
Rab5b
Map1lc3a
Gabarap
Map1lc3b
Snap29
Tecpr1
Epg5
Gabarapl2
Vamp8
Plekhm1
Stx17
Atg14
Autophagosome-lysosome fusion
Endosome-lysosome fusion
Shared tethering
log2FC (LPS vs. ctrl), paired t-test
