## Supplementary material for "Inflammatory proteostasis reprogramming creates a lysosomal bottleneck for microglial aggregate clearance": Figure S7

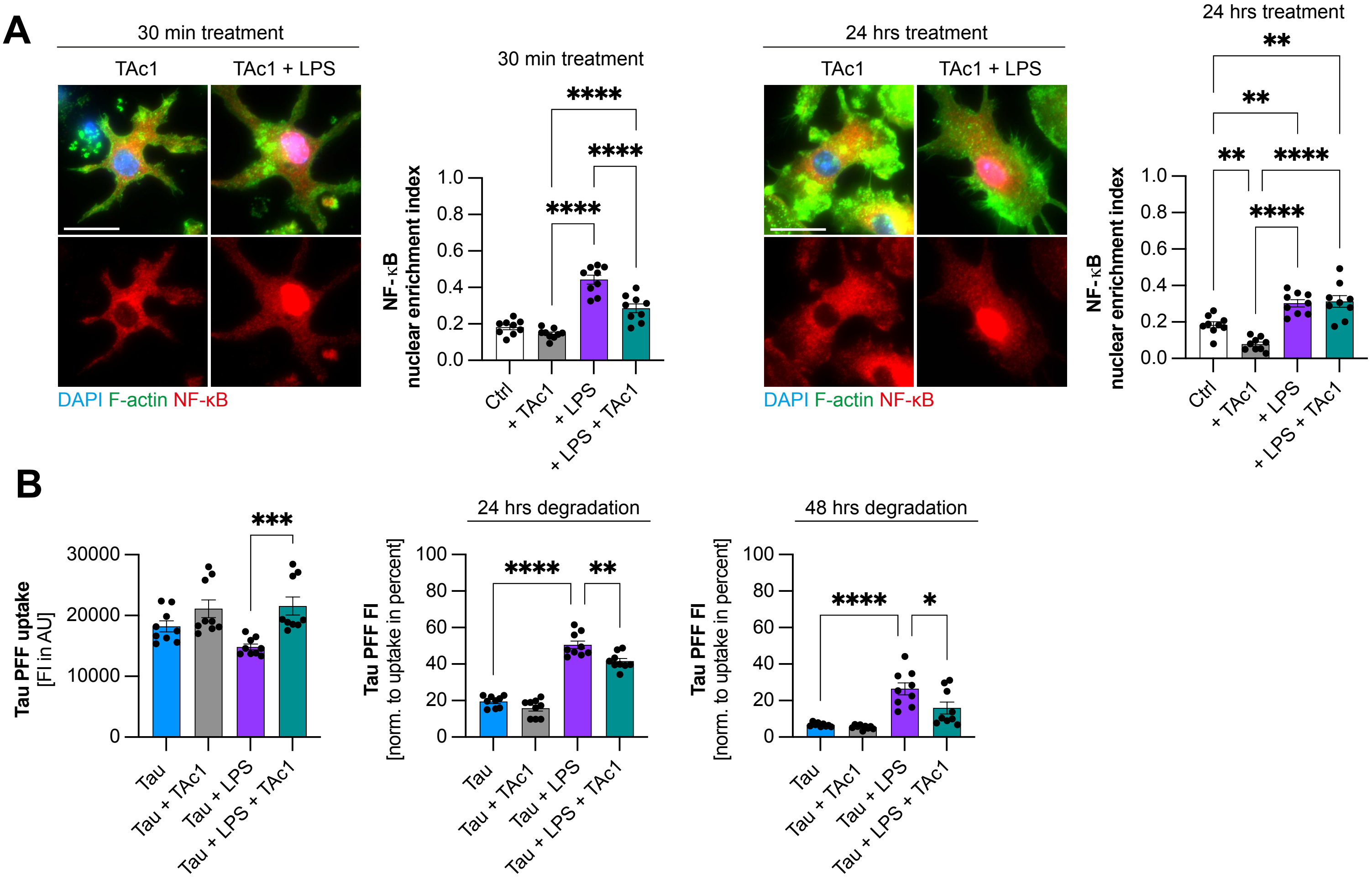
A
30 min treatment
TAc1
TAc1 + LPS
DAPI F-actin NF-κB
30 min treatment
NF-κB nuclear enrichment index
Ctrl
+ TAc1
+ LPS
+ LPS + TAc1
24 hrs treatment
TAc1
TAc1 + LPS
DAPI F-actin NF-κB
24 hrs treatment
NF-κB nuclear enrichment index
B
Tau PFF uptake [FI in AU]
Tau
Tau + TAc1
Tau + LPS
Tau + LPS + TAc1
24 hrs degradation
Tau PFF FI [norm. to uptake in percent]
48 hrs degradation
Tau PFF FI [norm. to uptake in percent]
